# Host species identity drives temporally stable octocoral microbiomes in Florida

**DOI:** 10.64898/2026.01.05.697751

**Authors:** Ronen Liberman, Matías Gómez-Corrales, Jose V. Lopez

## Abstract

Octocorals are foundational components of Caribbean reef ecosystems, yet their bacterial microbiomes remain poorly characterized compared to scleractinian corals. Understanding the relative contributions of host identity, morphology, and environmental factors in shaping these microbial partnerships is critical for predicting octocoral responses to environmental change. We characterized bacterial communities associated with four ecologically important Florida octocoral species over three years using 16S rRNA gene sequencing: the encrusting species *Briareum asbestinum* and *Erythropodium caribaeorum*, and the branching species *Eunicea flexuosa* and *Muricea muricata*. Host species identity emerged as the strongest predictor of microbiome composition, explaining 32.3% of bacterial community variation and substantially outweighing temporal and spatial effects. Core microbiome analysis revealed species-specific bacterial associations, with all four species harboring distinct *Endozoicomonas* variants as dominant symbionts. Encrusting species hosted taxonomically diverse microbiomes, while branching species maintained highly specialized communities dominated by *Endozoicomonas* and *Mycoplasma*. Bacterial communities within each host species remained stable across sites, depths, and years, with temporal and spatial factors explaining only 1.8% and 2% of variation, respectively. These findings demonstrate strong host control over microbial colonization and indicate that stable octocoral-bacteria associations may contribute to octocoral ecological resilience on western Atlantic reefs.

## Introduction

The ecological success of benthic coral reef organisms depends fundamentally on symbiotic partnerships with microorganisms (Rosenberg *et al*. 2007; Bourne, Morrow and Webster 2016; Van Oppen and Blackall 2019). Over the past two decades, advances in molecular and computational approaches have transformed our appreciation of these associations, establishing bacterial symbionts as essential contributors to host health, function, and adaptation. These bacterial communities, along with archaea, viruses, fungi, and unicellular protists, form an integral component of the coral holobiont (Rosenberg *et al*. 2007), the collective term for the host organism and all its diverse microbial associates. Within this holobiont framework, bacteria perform critical functions including nutrient cycling, pathogen defense, and metabolic support (Reshef et al., 2006; Bourne et al., 2016). As coral reef ecosystems face unprecedented environmental pressures, understanding the dynamics and stability of these bacterial partnerships has become both scientifically compelling and ecologically urgent (Voolstra *et al*. 2024).

A central question in coral symbiosis research concerns the degree to which coral microbiomes are shaped by intrinsic host properties versus external environmental factors (Pollock *et al*. 2018). At one extreme, certain bacterial symbionts may co-evolve with their coral hosts, becoming essentially heritable entities that remain stable across environmental gradients (Bordenstein & Theis, 2015; Rosenberg & Zilber-Rosenberg, 2018). Conversely, corals may dynamically acquire bacterial partners that enhance performance under specific conditions, with microbial communities shifting in response to environmental change (Webster & Reusch, 2017; Ziegler et al., 2017; Osman et al., 2020). Understanding where along this continuum coral-bacterial associations fall is critical for predicting how these foundation species will respond to rapidly changing ocean conditions.

The concept of a coral ’core microbiome’ has emerged as a useful framework for understanding host-microbe specificity and stability. Core microbiomes consist of bacterial taxa that are consistently associated with a coral species across spatial and temporal scales (Ainsworth *et al*. 2015; Hernandez-Agreda, Gates and Ainsworth 2017; Van Oppen and Blackall 2019). While definitions vary, core microbiomes generally consist of a reduced number of bacterial taxa that occur at low relative abundances but exhibit high prevalence across individuals (Hernandez-Agreda et al., 2018). These core taxa often exhibit phylosymbiosis, whereby bacterial diversity patterns mirror host phylogenetic relationships, suggesting long-term co-evolutionary associations (Pollock et al., 2018). Among coral-associated bacteria, members of the genus *Endozoicomonas* have emerged as particularly widespread and abundant core symbionts across diverse marine hosts, particularly cnidarians, including both hard (scleractinian) and soft (octocorals) corals (Neave et al., 2016; van Oppen & Blackall, 2019).

Octocorals (class Anthozoa, subclass Octocorallia) represent foundational yet understudied components of coral reef ecosystems. These sessile colonial cnidarians—commonly known as soft corals and gorgonians— possess eight-fold symmetry in their polyps and their internal skeletons is composed of microscopic sclerites rather than the massive calcium carbonate structures built by their hard coral relatives. Yet, octocorals still provide an essential three-dimensional habitat framework, particularly on Caribbean reefs where scleractinian coral cover has declined precipitously (Lenz *et al*. 2015; Lasker *et al*. 2020; Jones *et al*. 2022). Despite their ecological importance as habitat engineers, octocoral microbiomes remain far less characterized than those of reef-building scleractinians, with most studies focusing on a limited number of species or geographic regions (van de Water et al., 2018a).

Recent investigations of octocoral microbiomes have revealed patterns consistent with findings from scleractinian corals. Host species identity typically emerges as the strongest determinant of bacterial community composition, with each octocoral species harboring a distinctive microbial signature (Correa et al., 2013; van de Water et al., 2016; Reigel & Hellberg, 2023). Temperate Mediterranean gorgonians exhibit stable core microbiomes dominated by *Endozoicomonas* and other specific bacterial lineages that persist across seasons and geographic locations (van de Water et al., 2018b). Similar patterns have been documented in tropical octocorals from the Caribbean and Indo-Pacific, where host-specific bacterial assemblages show remarkable temporal stability despite strong environmental fluctuations (McCauley et al., 2020; Monti et al., 2023; Reigel & Hellberg, 2023). However, the extent to which octocoral microbiomes can shift in response to environmental variation may depend on host genetic structure, with genetically differentiated populations potentially showing greater microbiome plasticity than those of genetically homogeneous ones (Reigel & Hellberg, 2023).

Additional to species identity, host morphology represents another potentially important driver of microbial community structure. Octocorals display diverse growth forms, from encrusting mats that spread across hard substrates to erect branching colonies extending into the water column. These morphological differences may influence microbiome assembly through differential exposure to environmental microbes. Encrusting species, which maintain direct contact with reef substrata, experience microbial recruitment from both benthic surfaces and the water column, potentially resulting in more diverse bacterial communities (McCauley et al., 2016). By contrast, branching species recruit microbes primarily from the water column, which may constrain community diversity. Such morphology-dependent patterns have been observed in other benthic marine taxa, including sponges and seaweeds (Gloeckner et al., 2014; Lemay et al., 2021), but remain largely underexplored in octocorals.

The Florida Reef Tract represents an ideal system for investigating octocoral microbiome ecology. This reef system, which extends from the Dry Tortugas through the Florida Keys to southeastern Florida, harbors diverse octocoral assemblages spanning multiple morphological and phylogenetic groups (Goldberg, 1973). Unlike many Caribbean reefs where scleractinian corals have ecologically collapsed, Florida’s reefs maintain abundant and healthy octocoral populations that provide critical ecosystem services (Ruzicka et al., 2013, Liberman et al. 2025). However, these reefs face multiple stressors, including warming ocean temperatures, disease outbreaks, and coastal development, making it imperative to understand the stability and potential flexibility of octocoral-microbe associations. Previous microbiome studies from this region have been limited in taxonomic and temporal scope, leaving fundamental questions about microbiome drivers and stability unresolved.

Here, we characterize the bacterial communities associated with four ecologically important and widespread octocorals from southeastern Florida: the encrusting species *Briareum asbestinum* (Briareidae) and *Erythropodium caribaeorum* (Anthothelidae), and the branching species *Eunicea flexuosa* and *Muricea muricata* (Plexauridae). Through multi-year sampling across multiple reef sites and depths, we investigate the relative influences of host species identity, morphology, spatial location, and temporal variation on bacterial community composition. We investigate whether these octocorals harbor species-specific core microbiomes, examine the prevalence and diversity of *Endozoicomonas* across hosts, and assess whether host morphology constrains microbial diversity and community structure. By examining these factors simultaneously within a common ecological context, this study provides new insights into the drivers of octocoral microbiome assembly and stability in the western Atlantic, with implications for understanding how these foundational species may respond to ongoing environmental change.

## Methods

### Octocoral and seawater collection

In order to examine the octocoral bacterial assemblages in Florida (USA), specimens of *B. asbestinum, E. caribaeorum, E. flexuosa,* and *M. muricata* were collected by SCUBA from four locations (see supplementary Table S1) in three sampling campaigns: July-September 2023 and 2024, and January-February 2025.

The samples were collected from two inner reef sites (as defined by Walker, 2012), at a depth of 5-8 m, and two middle reef sites at a depth of 15-20 m (Figure 1). The collections were performed under a Florida Fish and Wildlife Conservation Commission Fishing License and a Special Activity License (24-2666-SRP). The octocorals host species were first assigned to species in the field based on their morphology. In each location, 4-5 specimens of each species were collected. Individuals were cut with scissors, placed in individual Whirl Pak bags filled with ambient seawater and brought to the surface. Samples were immediately frozen on dry ice and transported back to the laboratory (within 1-5 hrs of collection). Subsequently, all samples were rinsed off with filtered seawater (0.22 μm) and stored at −80°C for long-term storage. Seawater was also collected each time from the dive site in triplicates. These seawater samples (0.5 L) were used as environmental controls, to confirm that microbial communities associated with the octocorals were specific to them and not amplified from surrounding seawater DNA. Seawater (0.5 L) samples were filtered in 0.22 µm Sterivex cartridges (Millipore, SVGP01050), frozen and stored at −80 °C until processed.

**Figure 1.**
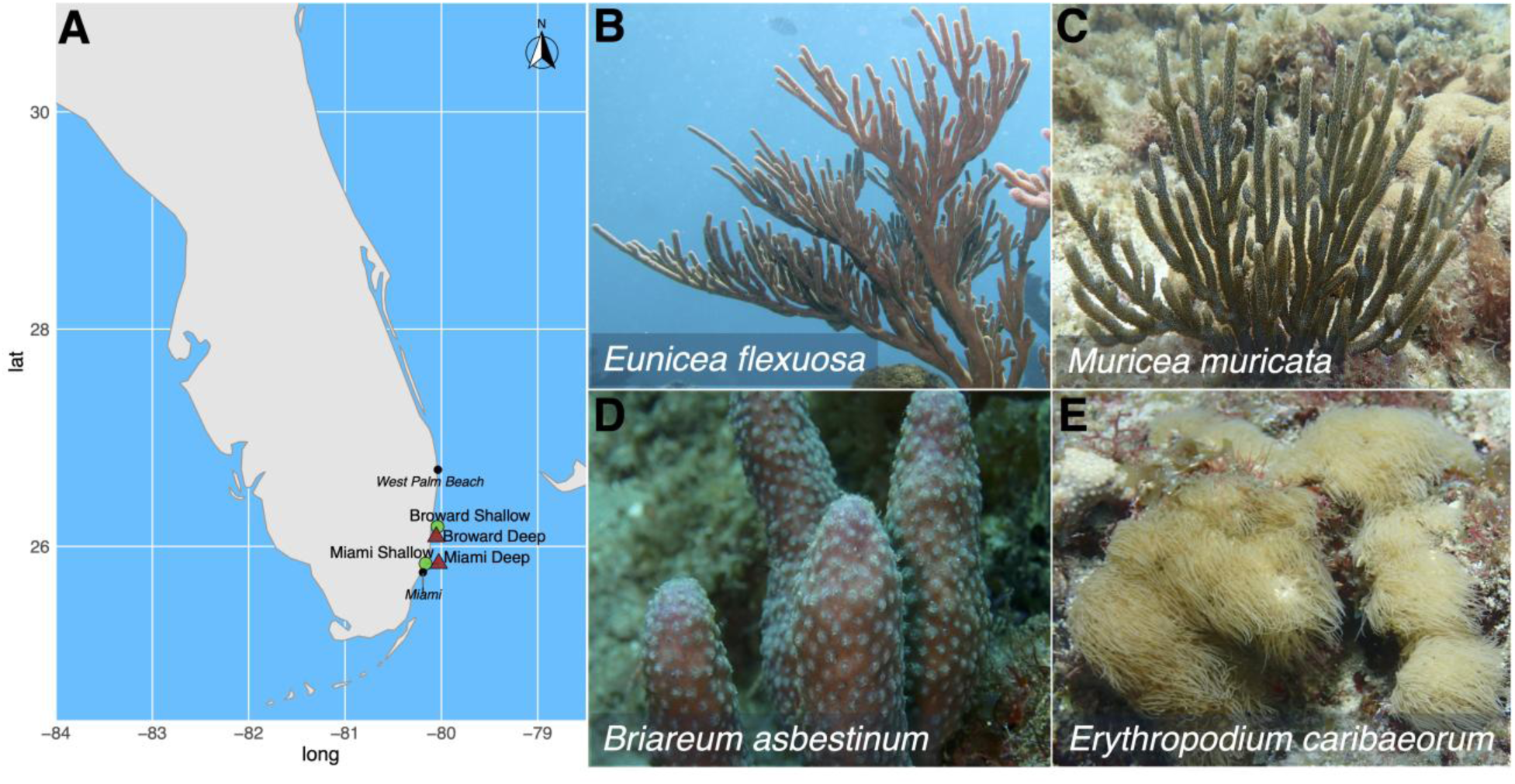
Study sites and octocoral species sampled from southeastern Florida reefs. (A) Map of collection locations in Miami-Dade and Broward counties showing four sampling sites: Miami Shallow (5-8 m depth), Miami Deep (15-18 m), Broward Shallow (5-8 m), and Broward Deep (15-20 m). Photographs show representative colonies of the four octocoral species examined: *(B) Eunicea flexuosa, (C) Muricea muricata, (D) Briareum asbestinum,* and *(E) Erythropodium caribaeorum.* Samples were collected between July 2023 and February 2025.

### DNA Extraction

Octocoral and seawater DNA were extracted using the Qiagen Powerlyzer PowerSoil DNA Extraction kit (Qiagen). Briefly, 2-5 polyps if each sample were excised using a sterile scalpel razor and then transferred into a Qiagen powerbead tube and resuspended in 750μl of lysis buffer, 60 μl of C1 and 25 μl of proteinase-K (20 mg/ml) following methods by Sunagawa et al. (2010) and Reigel and Hellberg (2023). These octocoral samples were incubated overnight at 65 °C (16-19 hrs.). Afterwards, a mechanical lysis was performed using a bead beating step (MoBIO, 3 X 30 seconds rounds at 6 m/sec). The samples were then centrifuged for 3 min at 10,000 *g*. The supernatant was transferred into a new tube and then subjected to the standard protocol of the Dneasy Powerlyzer PowerSoil kit (Qiagen) as per EarthMicrobiome standards (EMP; www.earthmicro biome.org; Thompson et al, 2017). DNA from the Sterivex filters were also extracted with the Qiagen PowerSoil kit to avoid yield discrepancy between DNA extraction protocols. The filters were placed into bead tubes (provided by the kit) and cut into fine pieces using sterile dissection scissors. Seawater DNA was extracted according to the manufacturer’s instructions using a 1 min bead-beating step.

### Octocoral host identification

To confirm the taxonomic identity of samples that deviated from observed species clustering patterns inferred from the microbiome data, we amplified the mitochondrial gene *mtMuts* using PCR. The gene was amplified using the *ND42599F* (5’ GCCATTATGGTTAACTAT TAC 3’) and *MUT3458R* (5’ TSGAGCAAAAGCCACTCC 3’) primers (Sánchez *et al*. 2003). PCR reactions were carried out in 20 μL volumes containing 1 μL of 1:10–1:40 diluted genomic DNA, 10 μL of Platinum™ Hot Start PCR 2× Master Mix (Thermo Fisher Scientific), 0.5 μL of each primer (10 μM), and 8 μL of nuclease-free water. The thermal cycling profile consisted of an initial denaturation at 94°C for 2 min; 48 cycles of 94°C for 90 s, 48°C for 90 s, and 72°C for 60 s; followed by a final extension at 72°C for 5 min. PCR products were visualized on 1.5% agarose gels stained with GelRed. Amplicons were purified in PCR-grade water and quantified using a Qubit 2.0 Fluorometer (Invitrogen). Bidirectional Sanger sequencing was performed by Azenta Life Sciences (South Plainfield, NJ, USA). Resulting chromatograms were inspected and edited in 4Peaks v1.8 (Nucleobytes, 2004), and sequences were aligned in AliView v1.3 (Larsson 2014) using MAFFT v7 (Katoh and Standley 2013). We identified the closest matching sequence for each amplicon using BLASTn searches against the NCBI GenBank non-redundant nucleotide database and recorded sequence identity and query coverage percentages for each match.

We constructed *mtMuts* phylogenetic trees for each species using IQTREE3 v3.0.1 with automatic model selection under the ModelFinder algorithm and 1,000 ultrafast bootstrap replicates to assess node support (Nguyen *et al*. 2015; Kalyaanamoorthy *et al*. 2017; Hoang *et al*. 2018). For each species, additional *mtMutS* sequences and their closest BLASTn matches were downloaded from NCBI to serve as outgroups for tree rooting. Resulting consensus trees were imported into R and rooted using the **“**ape” package (Paradis and Schliep 2019). Tree visualization and annotation were performed using the “ggtree” R package (Yu *et al*. 2017).

### 16S rRNA gene amplicon sequencing and analysis

The 16S rRNA gene was amplified using standard 515F and 806R universal primers (Caporaso et al. 2011, Apprill et al. 2015). This primer set was chosen because it targets a broad range of bacterial and archaeal taxa with the exception of only a few groups (Bates et al., 2011; Caporaso et al., 2011). The primers amplified ∼300 bp fragments spanning the gene’s V3 and V4 regions. PCR products were visualized on 1.5% agarose gel. Successful reactions (i.e., with a clear single band) were then amplified using barcoded 515F and 806R primers, with each sample receiving identical barcoded 806R primer sequences and unique barcoded 515F primer sequences. A final 1.5% agarose gel was run to confirm the successful barcoding of the samples and successful reactions were quantified using a Qubit 4.0 fluorometer (Life Technologies). Afterwards samples were diluted to 4nM using nuclease-free water to ensure that each was represented in equimolar amounts. Diluted samples were then pooled together, cleaned using AMPure XP beads (Beckman Coulter, Beverly, MA), and checked for quality and contamination using the Agilent TapeStation 4150. The final library pool was then loaded into a MiSeq instrument (Illumina Inc), using the MiSeq Reagent Kit v3 at 600 cycles following the EMP standard protocols (Thompson *et al*. 2017).

### Bioinformatic analyses

DNA sequences were run through Quantitative Insights into Microbial Ecology v2023.2 (QIIME2) pipeline. The demultiplexed forward and reverse sequence reads were run for quality filter, assign taxonomy to amplicon sequence variants (ASVs), reconstruct their phylogeny, and produce abundance analysis. Quality filtering and data trimming were conducted with the ‘qiime dada2 denoise-paired’ command. The QIIME2-generated sequences were assigned to ASVs at 99% sequence identity using the SILVA classifier (v138.2). Reads identified as mitochondria, chloroplast and/or host contaminants were removed before any statistical analysis.

#### Statistical Analyses

All diversity analyses were conducted on rarefied ASV abundance data to normalize for sequencing depth differences across samples, following established protocols for microbiome studies (Weiss et al., 2017). ASV tables were rarefied to 20,000 reads per sample using QIIME2, determined by rarefaction curve analysis (Supplementary Figure 1). This rarefaction depth balanced adequate sampling coverage while minimizing sample loss due to low read counts. This ultimately led to the removal of four octocoral samples that did not meet the above-mentioned standards.

Alpha diversity metrics (observed ASV richness, Shannon diversity, and inverse Simpson diversity indexes) were calculated from the rarefied data using the “microeco” R package (Liu et al., 2021). Data normality was assessed using Shapiro-Wilk tests (α = 0.05) (R Core Team, 2023). Kruskal-Wallis tests evaluated overall differences among the four octocoral species using the “stats” package, followed by Dunn’s post-hoc tests with Bonferroni correction for multiple pairwise comparisons using the FSA package (Ogle et al., 2023). Wilcoxon rank-sum tests assessed temporal changes within each species across the three-year sampling period.

For beta diversity analysis, Bray-Curtis dissimilarity matrices were calculated from the rarefied data using the “vegan” package (Oksanen et al., 2022), and Principal coordinate analysis (PCoA) was used to visualize community composition patterns. PERMDISP tests (999 permutations) verified homogeneity of group dispersions prior to PERMANOVA analyses. PERMANOVA was conducted following a hierarchical approach. Primary analysis tested overall species effects across all samples. Nested analyses examined temporal effects (year) and spatial effects (site) within each octocoral species separately. Multiple pairwise comparisons between all species combinations and between octocorals versus seawater controls were performed with Bonferroni correction. F-statistics, R² values (proportion of variance explained), and p-values were calculated for all tests. Effect sizes were interpreted as: R² >0.25 (large), 0.09-0.25 (medium), <0.09 (small). Statistical significance was set at α = 0.05.

Core bacterial communities were identified following Van Oppen & Blackall (2019), using multiple prevalence thresholds. ASVs were classified as core members if present in ≥75% (moderate), ≥90% (stringent), or 100% (sensu stricto) of samples within each octocoral species. All analyses were conducted at the ASV level. To ensure robustness of the 90% core microbiome against sampling variability, we implemented bootstrap resampling validation. For each species, 1,000 bootstrap iterations were performed, with each iteration randomly sampling 20 individuals with replacement from the total sample pool for that species. In each bootstrap iteration, ASVs meeting the 90% prevalence threshold were identified. ASVs were retained in the final validated 90% core microbiome only if they appeared as core members in ≥80% of bootstrap iterations, ensuring statistical robustness while accounting for technical and biological variation. This bootstrap approach provides confidence that identified core members represent truly stable associations rather than artifacts of sampling or stochastic variation. Bootstrap frequency percentages are reported for all validated core ASVs, with higher frequencies indicating greater stability across sampling scenarios (see supplementary material). For all prevalence thresholds, prevalence was calculated as the percentage of samples within each species containing each ASV (presence/absence), and mean relative abundance was calculated as the average proportional abundance of each core ASV across all samples within that species.

ANCOM-BC2 identified bacterial taxa with significantly different abundances among octocoral species using the ‘ANCOMBC’ package (Lin and Peddada 2020). The analysis incorporated bias correction for uneven sampling and library composition differences. False discovery rate (FDR) correction controlled for multiple testing across ASVs. Significant taxa were defined as those with FDR-corrected p < 0.05.

## Results

### Sequencing results

16S rRNA gene sequencing generated a total of 18,924,015 high-quality reads across all samples after quality filtering and removal of chloroplast and mitochondrial sequences (Supplementary Table S2). Reads were assigned to amplicon sequence variants (ASVs) across the entire dataset. The number of samples varied among octocoral species, with seawater controls also included for comparison.

### Octocoral host identification

Molecular identification of octocoral hosts that deviated from observed microbiome clustering patterns was confirmed through Sanger sequencing of the mitochondrial *mtMuts gene* and BLAST analysis against the NCBI GenBank database (Table 1). All samples across our four species exhibited high sequence identity (>98%) and query coverage (>90%), allowing us to confidently assign deviated samples to their true taxon. *B. asbestinum* (n = 17), *E. caribaeorum* (n = 23), *E. flexuosa* (n = 12), and *M. muricata* (n = 20) all showed sequence identity values ranging from 98.23% to 100.00%, validating the species assignments used throughout this study.

**Table 1.**
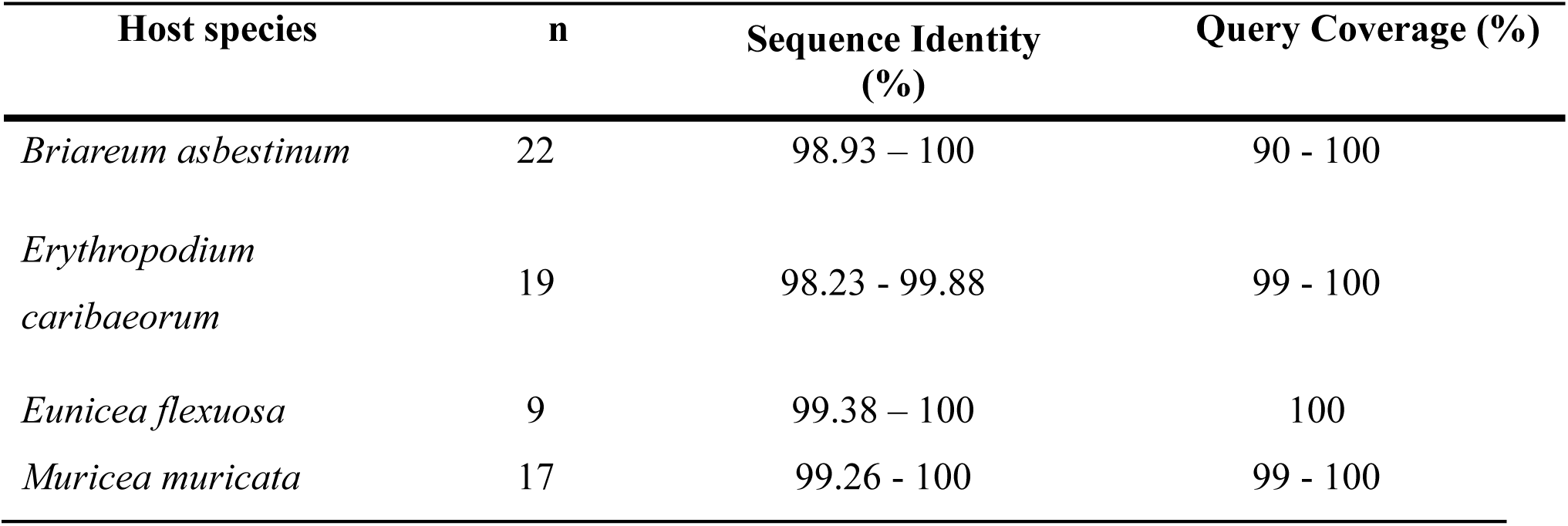
Host species identification summary based on Sanger sequencing of mitochondrial markers (*mtMuts*) and BLAST comparison with NCBI GenBank sequences. . Values represent the range of sequence identity and query coverage across all successfully sequenced samples for each species. n = number of samples per species.

### Bacterial community composition: host differentiation

The bacterial communities associated with the four octocoral species exhibited distinct taxonomic profiles at the phylum level (Figure 2). Proteobacteria dominated the microbiomes of *B. asbestinum, E. caribaeorum,* and *E. flexuosa,* while *M. muricata* was primarily dominated by Firmicutes. Notably, phylum-level diversity differed substantially among host species, with *E. caribaeorum* harboring the most diverse community (10 phyla contributing ≥1% mean relative abundance, collectively accounting for 94.81% of the total community), followed by *B. asbestinum* and *M. muricata* (6 phyla each, 94.37% and 98.04% respectively), and *E. flexuosa* (5 phyla, 95.90%). The composition of dominant phyla varied markedly among hosts: *E. caribaeorum* uniquely harbored substantial abundances of Actinobacteriota, Crenarchaeota, Acidobacteriota, Myxococcota, and NB1-j, while *M. muricata* was characterized by enrichment of Firmicutes, Spirochaetota, and Campilobacterota. Temporal shifts in community composition were apparent across the three-year sampling period (2023-2025), with the most pronounced changes observed in *E. caribaeorum*.

**Figure 2.**
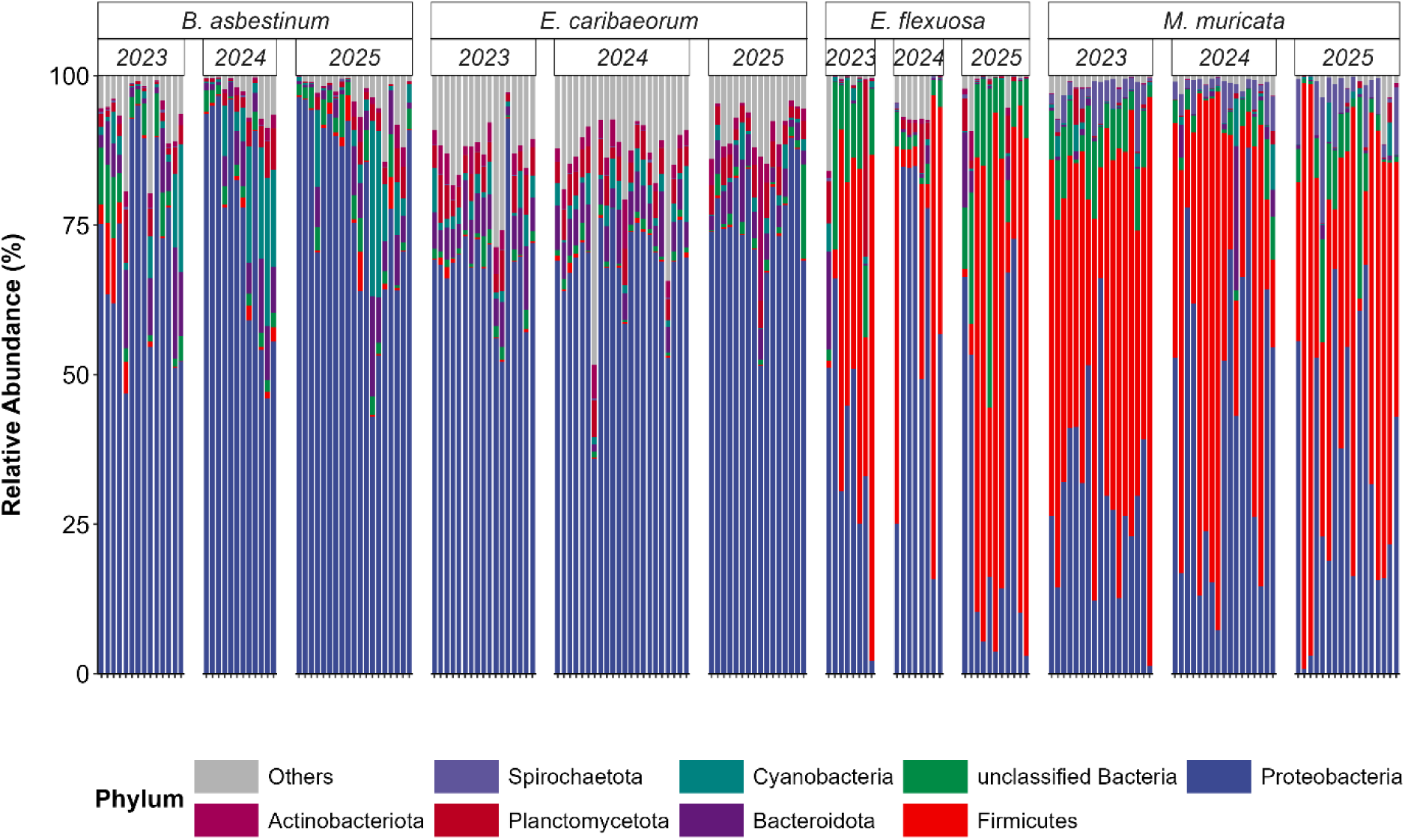
Phylum-level bacterial community composition in four octocoral species across sampling years. Stacked bar charts showing the relative abundance of the eight most abundant bacterial phyla in *Briareum asbestinum*, *Erythopodium caribaeorum*, *Eunicea flexuosa*, and *Muricea muricata* from 2023 and 2025. Sequences were rarefied to 20,000 reads per sample to normalize sequencing depth.

### Core octocoral bacterial community

To identify the most consistently associated bacterial taxa, we employed a multi-threshold presence-based approach to define core bacterial communities. ASVs were classified as core members if present in ≥75%, ≥90%, or 100% of samples within each octocoral species (Figure 3, Supplementary Table S3-4). This analysis revealed substantial differences in bacterial community complexity among the four octocoral species across all threshold levels. At the 90% prevalence threshold, core microbiome analysis revealed a wide range of ASVs, spanning from a single ASV to 24 ASVs. *E. caribaeorum* exhibited the most taxonomically diverse core microbiome, harboring 24 microbial ASVs representing multiple families including Endozoicomonadaceae, Rhodobacteraceae, Woeseiaceae, Spongiibacteraceae, Hyphomicrobiaceae, and Hyphomonadaceae. Notably, this species’ core also included members of the archaeal genus *Candidatus Nitrosopumilus*, suggesting potential involvement in nitrogen cycling processes. *B. asbestinum* maintained a moderately sized core of 6 ASVs, dominated by members of Endozoicomonadaceae and Rhodobacteraceae families. In contrast, the branching species *M. muricata* and *E. flexuosa* exhibited remarkably streamlined cores, with only 2 and 1 ASVs, respectively.

**Figure 3.**
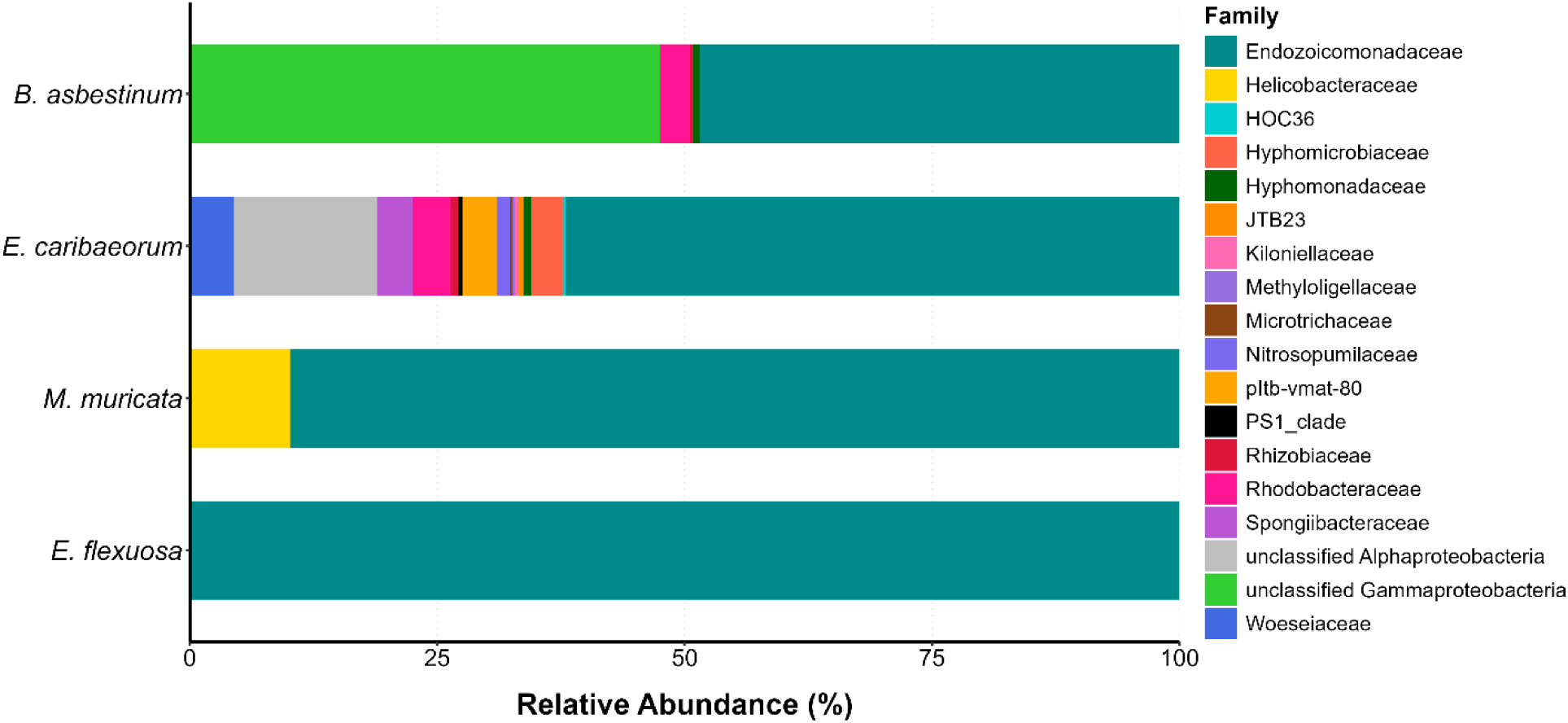
Relative abundance of bacterial families in the core microbiome of four octocoral species. Stacked horizontal bars show the proportional contribution of each bacterial family to the core microbiome, defined as ASVs present in ≥90% of samples within each species and validated through bootstrap resampling (1,000 iterations 20 randomly selected samples per iteration). Each bar represents 100% of the core microbiome composition for that species.

*Endozoicomonas* dominated the 90% core microbiome across all four species, with species-specific variants present in *B. asbestinum* (2 ASVs, combined 18.55% mean relative abundance), *E. caribaeorum* (3 ASVs, combined 19.16%), *M. muricata* (1 ASV, 7.72%), and *E. flexuosa* (1 ASV, 7.42%). Notably, *B. asbestinum* was the only species where *Endozoicomonas* was not the most abundant core member at this threshold; instead, an unclassified Gammaproteobacteria ASV dominated the core (18.18%), indicating a potential different microbial association strategy in *B. asbestinum* compared to other octocorals examined.

Additional species-specific patterns emerged in core microbiome composition. The Rhodobacteraceae family was represented in both encrusting species’ cores (*B. asbestinum*: 1.16%; *E. caribaeorum*: 1.17%), suggesting this group may play an important role in encrusting octocoral-bacterial associations. *E. caribaeorum* uniquely harbored two *Filomicrobium* ASVs (combined 1 % abundance) and members of the archaeal genus *Candidatus Nitrosopumilus* (0.39%), potentially contributing to nitrogen transformation in this species. *M. muricata*’s core included a member of Helicobacteraceae (0.87%), a family not detected in the cores of other species. Additional taxa present in *E. caribaeorum*’s diverse core included members of Woeseiaceae (1.37%), Spongiibacteraceae (1.11% combined), and multiple unclassified Alphaproteobacteria lineages, reflecting the complex bacterial assemblage associated with this encrusting species. In contrast, *E. flexuosa* maintained an exclusive *Endozoicomonas* association with no additional taxa at 90% prevalence.

Comparisons across prevalence thresholds revealed that encrusting species maintained substantially larger cores than branching species at all thresholds examined. *E. caribaeorum* exhibited the most dramatic reduction in core size, declining from 90 ASVs at 75% prevalence to 24 ASVs at 90%, and ultimately to 5 ASVs at the sensu stricto (100%) threshold. In contrast, branching species (*M. muricata* and *E. flexuosa*) demonstrated highly stable cores, with minimal reduction across thresholds: *M. muricata* retained 2 ASVs at 90% compared to 1 ASV at 100% prevalence, while *E. flexuosa* maintained a single ASV across both the 90% and 100% thresholds.

### Alpha and beta diversity metrices reveal species-specific bacterial richness

Alpha diversity metrics demonstrated significant variation in bacterial community richness and evenness among the four octocoral species (Table 2, Figure 4). Markedly, *E. caribaeorum* harbored the most diverse bacterial communities across all metrics, with the highest observed ASV richness (1344 ± 404 in 2023), Shannon diversity (5.16 ± 0.99 in 2023), and inverse Simpson indices (54.6 ± 55.5 in 2023). In contrast, *M. muricata* consistently maintained the lowest diversity values, with ASV richness ranging from only 131 ± 77 to 176 ± 142 across years, suggesting a more specialized and stable microbiome. Interestingly, temporal dynamics in alpha diversity differed substantially among host species. While *B. asbestinum* and *E. flexuosa* maintained relatively stable diversity patterns across years, *E. caribaeorum* exhibited significant temporal shifts in both Shannon diversity (p < 0.001) and inverse Simpson indexes (p < 0.01) between sampling years. The Shannon diversity of *E. caribaeorum* increased from 5.16 ± 0.99 in 2023 to 5.35 ± 0.44 in 2024, before declining to 4.49 ± 0.65 in 2025. Similarly, *M. muricata* showed remarkable stability in bacterial diversity across the three-year period, with no significant temporal changes detected in any diversity metric.

**Figure 4.**
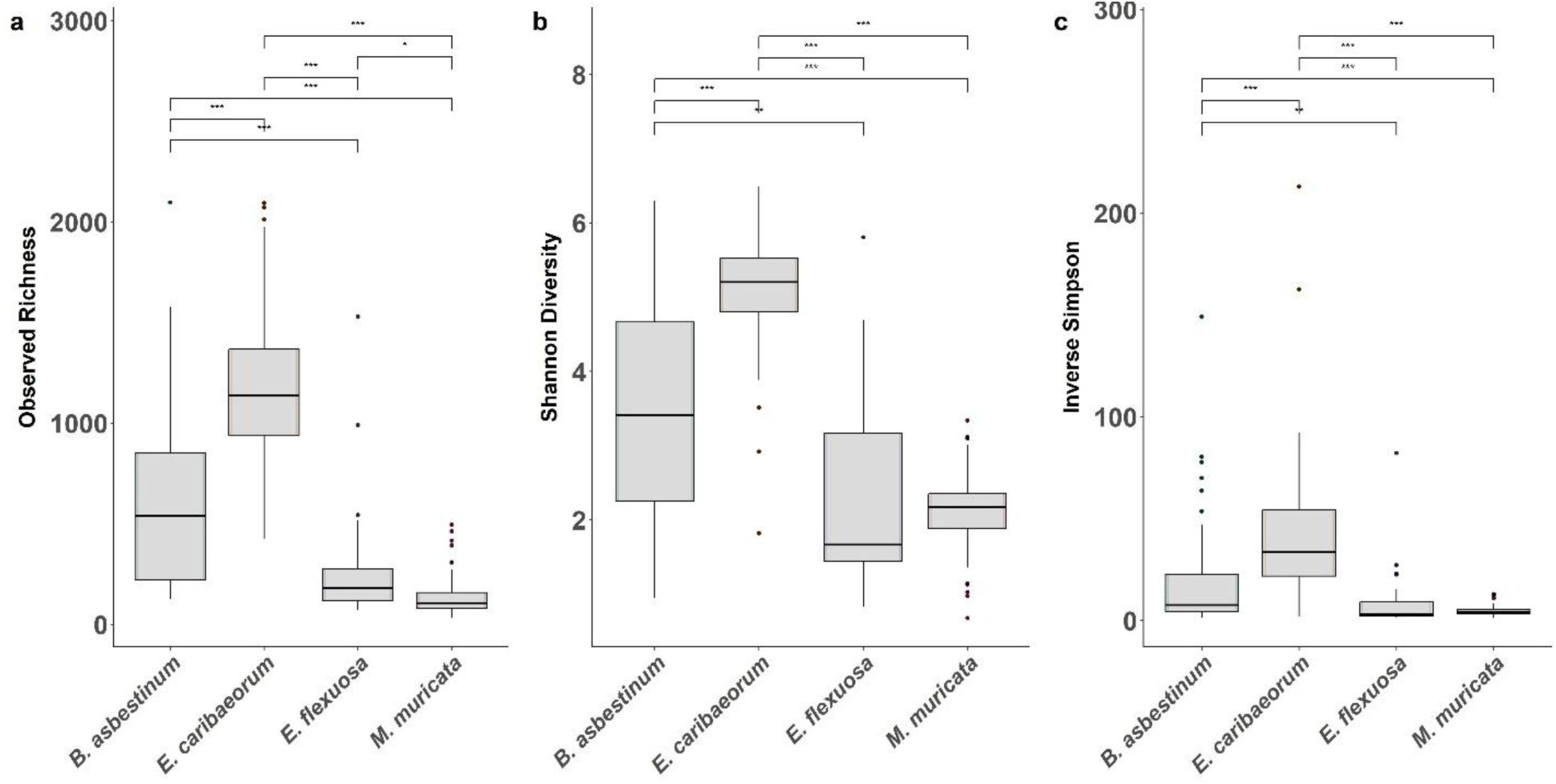
Alpha diversity metrics of bacterial communities across four octocoral species. Boxplots show (a) observed richness, (b) Shannon diversity index, and (c) inverse Simpson diversity index for bacterial communities associated with *Briareum asbestinum, Erythropodium caribaeorum, Muricea muricata, and Eunicea flexuosa.*. Data represents all samples collected from Broward and Miami counties from July 2023 and February 2025. Sequences were rarefied to 20,000 reads per sample to normalize sequencing depth. Box plots display median (center line), first and third quartiles (box boundaries), and 1.5× interquartile range (whiskers), with outliers shown as individual points. Significant differences among species were observed for all three diversity metrics (Kruskal-Wallis test, p < 0.05; see supplementary Table 6 for pairwise comparisons).

**Table 2.** Alpha diversity metrics of bacterial communities across four octocoral species and sampling years. Values represent mean ± standard deviation for ASVs (Amplicon Sequence Variants), observed richness, Shannon diversity index, and inverse Simpson diversity index for bacterial communities associated with *Briareum asbestinum, Erythropodium caribaeorum, Muricea muricata,* and *Eunicea flexuosa*. Data represents samples collected from Broward and Miami counties sampling locations between July 2023 and February 2025. Sequences were rarefied to 20,000 reads per sample to normalize sequencing depth (Kruskal-Wallis test: * p < 0.05, ** p < 0.01, *** p < 0.001).

| Species | Year | N | ASVs | Shannon Diversity | Inverse Simpson |
| --- | --- | --- | --- | --- | --- |
| *B. asbestinum* | 2023 | 14 | 802 ± 570 | 3.96 ± 1.36 | 32.5 ± 42.1 |
| | 2024 | 12 | 546 ± 362 | 3.46 ± 1.52 | 18.4 ± 22.6 |
| | 2025 | 19 | 515 ± 394 | 3.26 ± 1.32 | 13.8 ± 16.3 |
| *E. caribaeorum* | 2023 | 17 | 1344 ± 404 | 5.16 ± 0.99 | 54.6 ± 55.5 |
| | 2024 | 22 | 1242 ± 381 | 5.35 ± 0.44*** | 50.7 ± 23** |
| | 2025 | 16 | 919 ± 219 | 4.49 ± 0.65*** | 22.3 ± 14.5** |
| *E. flexuosa* | 2023 | 8 | 326 ± 490 | 2.4 ± 1.52*** | 14 ± 27.8** |
| | 2024 | 8 | 234 ± 144 | 2.65 ± 0.96 | 7.1 ± 4.4 |
| | 2025 | 11 | 278 ± 288 | 2.25 ± 1.48 | 8.7 ± 10.4 |
| *M. muricata* | 2023 | 17 | 176 ± 142 | 2.25 ± 0.52 | 5.1 ± 1.8 |
| | 2024 | 17 | 131 ± 77 | 2.06 ± 0.63 | 5.3 ± 3.5 |
| | 2025 | 17 | 137 ± 95 | 2.07 ± 0.57 | 4.5 ± 2.5 |

Principal coordinate analysis (PCoA) based on Bray-Curtis dissimilarity revealed clear separation of bacterial communities among octocoral species and between octocorals and seawater controls (Figure 5). The four octocoral species formed clearly distinct clusters with minimal overlap, indicating strong host-specific bacterial assemblages. Notably, the first two principal coordinates explained a substantial proportion of the variation in community composition (24.4%), with samples clustering tightly by host species regardless of collection year or site.

**Figure 5.**
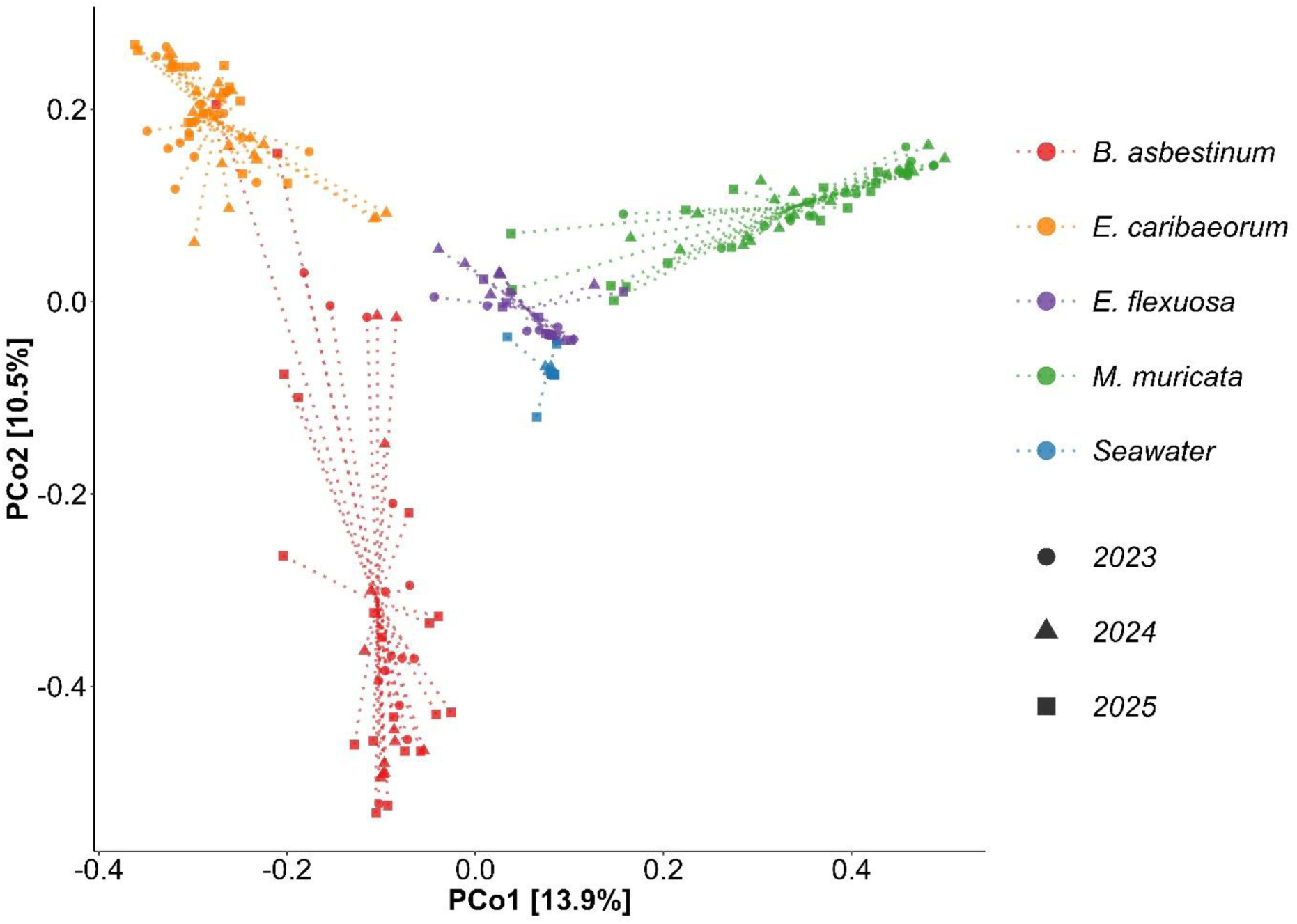
Principal coordinate analysis (PCoA) of octocoral-associated bacterial communities based on Bray-Curtis dissimilarity. Samples are colored by host species (red=*Briareum asbestinum, orange=Erythopodium caribaeorum, green=Muricea muricata,* and *purple=Eunicea flexuosa)* and seawater controls in blue, with shapes representing collection years (2023-2025). Each point represents the bacterial community composition of an individual sample, while centroids indicate the average community structure for each species. Samples were collected from shallow (5- 8 m) and deep (15-20 m) sites at Broward and Miami counties, Florida.

PERMANOVA analysis confirmed that host species identity was the strongest driver of bacterial community structure, explaining 32.3% of the total variation (F = 22.436, p < 0.001; Table 3). While temporal (year) and spatial (site) factors also significantly influenced community composition, their effects were markedly smaller, accounting for only 1.8% and 2.0% of the variation, respectively (Supplementary Table S5). This hierarchical pattern of variation underscores the dominance of host-specific factors in shaping octocoral microbiomes.

**Table 3.**
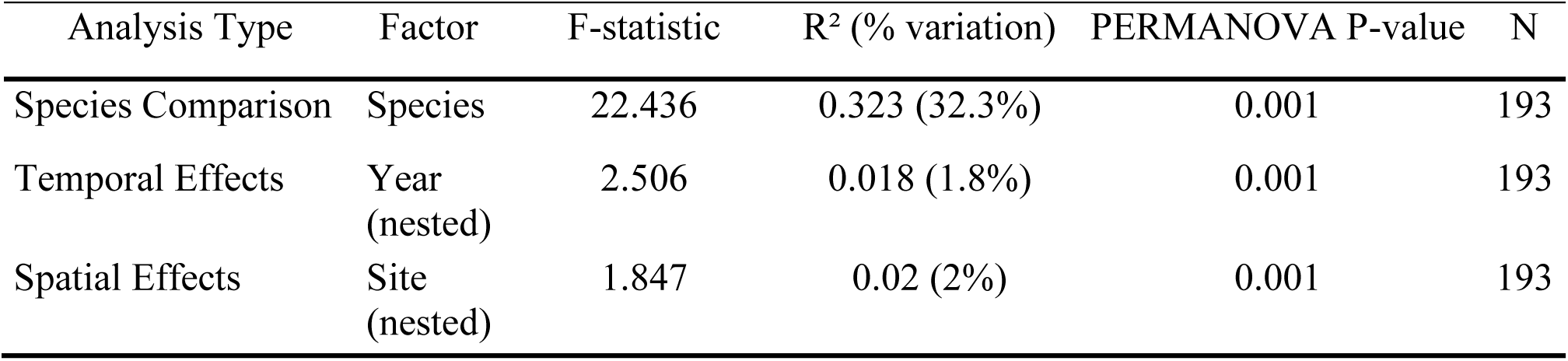
PERMANOVA results for factors affecting octocoral-associated bacterial community composition. . Analysis based on Bray-Curtis dissimilarity matrices showing the relative contribution of host species identity, temporal variation (year), and spatial variation (site) to bacterial community structure. F-statistics represent the strength of each factor’s effect, R² values indicate the proportion of total variance explained by each factor. Analysis included 193 samples from four octocoral species (*Briareum asbestinum, Erythopodium caribaeorum, Eunicea flexuosa, and Muricea muricata*) collected across multiple years (2023-2025) and sites in Florida. Sequences were rarefied to 20,000 reads per sample prior to analysis.

Pairwise comparisons revealed that all octocoral species harbored significantly different bacterial communities from each other and from seawater controls (Supplementary Table S6). Notably, the strongest differentiation occurred between *E. caribaeorum* and *M. muricata* (F = 35.056, R² = 0.252, p < 0.001), while *E. flexuosa* and *M. muricata* showed the least differentiation among octocoral pairs (F = 12.953, R² = 0.146, p < 0.001), although still highly significant. All octocoral species were strongly differentiated from seawater communities, with F-statistics ranging from 13.432 to 27.525 (all p < 0.001 after Bonferroni correction), confirming that octocoral-associated bacterial communities are distinct from the surrounding seawater microbiome.

### Temporal and spatial patterns among species

Individual species showed varying degrees of temporal and spatial structuring in their bacterial communities (Figures S5-7). Temporal effects were most pronounced in *M. muricata* (F = 2.120, p = 0.002, R² = 0.081) and *B. asbestinum* (F = 1.920, p = 0.005, R² = 0.084), while spatial variation was highest in *B. asbestinum* (F = 2.554, p = 0.001, R² = 0.157) and *E. flexuosa* (F = 1.541, p = 0.029, R² = 0.167). Notably, despite these significant effects, the proportion of variance explained by temporal and spatial factors within species remained relatively low (R² < 0.17 for all comparisons), further emphasizing the overriding importance of host identity in structuring octocoral bacterial communities.

### Differentially abundant taxa distinguish host species

ANCOM-BC2 analysis identified multiple bacterial taxa that were differentially abundant among the four octocoral species (Figure 6). Importantly, certain bacterial genera showed strong host-specific associations, with some taxa being nearly exclusive to particular octocoral species. *E. caribaeorum* was characterized by high abundances of several unique bacterial taxa not found in the other species, consistent with its higher overall diversity. In contrast, *M. muricata* was dominated by a smaller set of bacterial genera that was present at significantly higher relative abundances than in other hosts. Interestingly, while some bacterial taxa were shared among multiple octocoral species, their relative abundances varied significantly, suggesting that host-specific factors modulate the successful associations with particular bacterial lineages. The most differentially abundant taxa showed fold-changes exceeding an order of magnitude between host species, highlighting the strong selective pressures exerted by different octocoral hosts on their associated bacterial communities.

**Figure 6.**
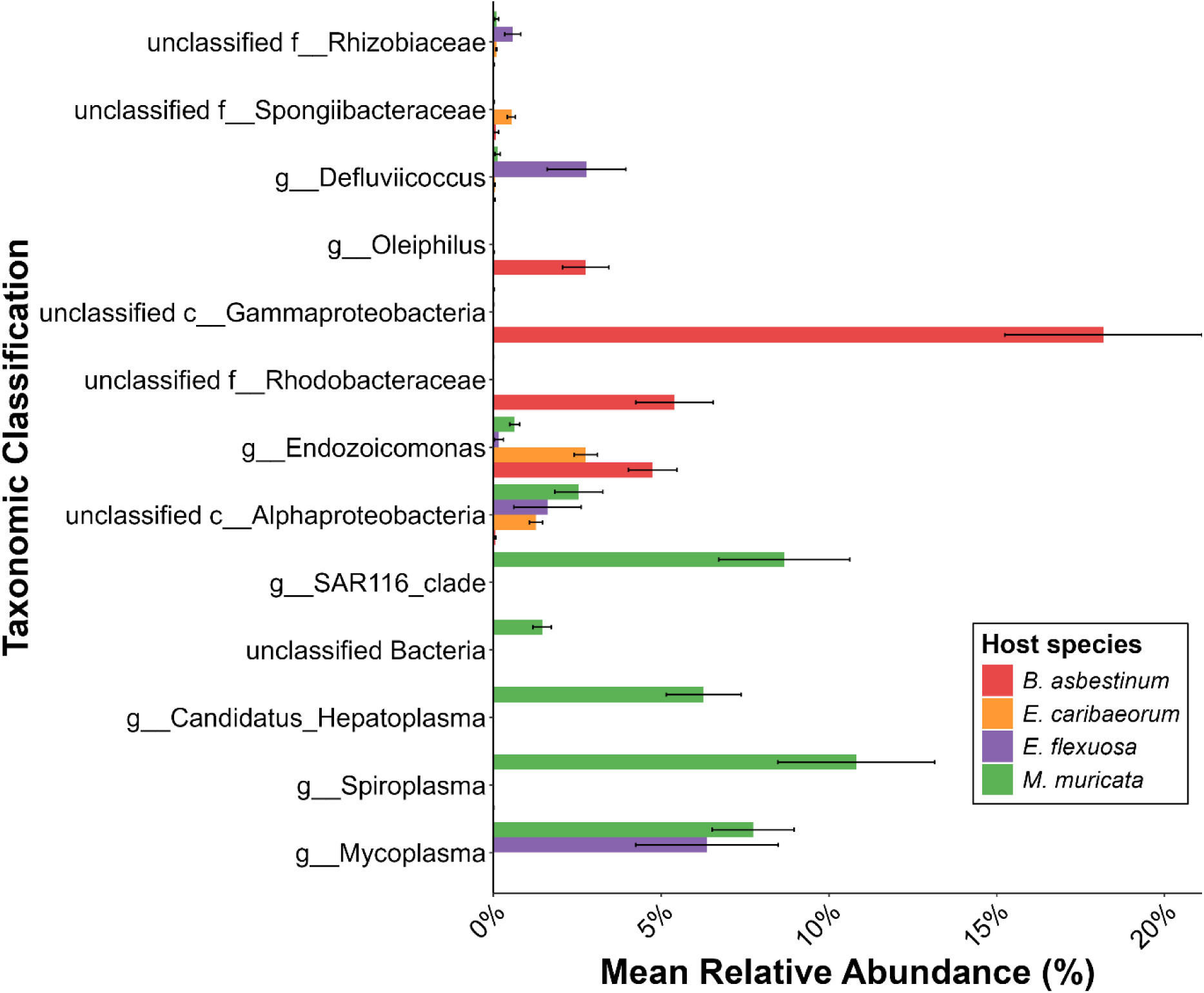
Differentially abundant bacterial taxa across four coral species. Mean relative abundance (± standard error) of bacterial taxa identified as significantly different between octocoral species using ANCOM-BC2 analysis. Only the most significant taxa (lowest adjusted p-values) are shown, representing *Briareum asbestinum* (red), *Erythopodium caribaeorum* (orange), *Eunicea flexuosa* (purple), and *Muricea muricata* (green). Taxonomic classifications represent the lowest resolved level, with unclassified groups labeled by their highest classified taxonomic rank. Error bars represent standard error of the mean. The displayed taxa showed significant differential abundance between all species (FDR-corrected p < 0.05).

## Discussion

### Octocoral host species harbor distinct microbiome communities

Host species identity emerged as the strongest determinant of bacterial community composition, explaining 32.3% of the total bacterial variation in our dataset (Table 3). Each host species harbored distinctive bacterial assemblages that remained consistent across sampling years and locations (Figure 5). While sample sizes varied among species, the consistent host-specific clustering across all sampling periods suggests these patterns are robust despite differences in statistical power. This pronounced host-specific clustering aligns with previous findings on western Atlantic and Caribbean octocorals, where each species maintained distinct bacterial communities (McCauley et al., 2020; Monti et al., 2023; Reigel & Hellberg, 2023).

Octocoral host phylogeny may also contribute to the observed microbiome differences, as observed in other taxa (Thomas *et al*. 2016; Moeller *et al*. 2023); however, it is difficult to disentangle this factor from species-level effects, as all four studied octocorals belong to different genera and three families within two different orders (McFadden et al., 2022). For example, it should be noted that *E. caribaeorum* and *B. asbestinum* represent two closely related taxa within the order Scleralcyonacea, compared to the more distantly related species *E. flexuosa* and *M. muricata,* in the order Malacalcyonacea. Yet, the former pair appear to have very divergent bacterial communities.

Additionally, differences in algal symbiont communities may influence octocoral-associated microbiomes (Lawson et al., 2018). Indeed, while all four species harbor photosynthetic dinoflagellates (family Symbiodiniaceae; LaJeunesse et al., 2018), these communities differ genetically: *E. caribaeorum* hosts algal symbionts of the genus *Cladocopium*, whereas *B. asbestinum*, *E. flexuosa*, and *M. muricata* are dominated by members of the genus *Breviolum* (Goulet and Coffroth 2004; Goulet *et al*. 2017). Symbiodiniaceae identity has been shown to influence bacterial community structure in cnidarian holobionts, with different algal genera associated with distinct bacterial assemblages (Röthig et al., 2016; McCauley et al., 2023), suggesting that the observed differences in Symbiodiniaceae composition among our octocoral species may contribute to their distinct bacterial communities. Therefore, the influence of host-specific effects suggests that intrinsic host factors, including tissue source, phylogeny, physiology, and associated symbionts play crucial roles in structuring octocoral, as well as many other animals, microbiomes.

### The influence of host morphology

Beyond species identity, host morphology emerged as a secondary but significant driver of microbiome composition, strongly influencing bacterial diversity levels. Microbiome diversity differed markedly between the two octocoral morphologies, which may reflect divergent ecological strategies and host-microbe relationships (Voolstra & Ziegler, 2020; Vohsen & Herrera, 2024). The encrusting species *E. caribaeorum* and *B. asbestinum* maintained the most diverse bacterial communities, potentially characterizing them as microbial generalists that foster more diverse and flexible microbial assemblages. This flexibility may confer adaptive advantages under variable environmental conditions by providing access to a broader range of microbial functions or ecological redundancy. Conversely, the two branching species, *E. flexuosa* and *M. muricata*, exhibited more specialized microbial communities characterized by lower α-diversity and reduced variability across temporal and spatial scales (Table 2, Figure 4). This pattern suggests a microbial specialist strategy, where these hosts maintain tight control over their bacterial associates, potentially optimizing specific host-microbe interactions at the expense of flexibility. This morphology-diversity relationship was further reinforced by the significant separation of encrusting and branching species in ordination analysis (Figure S8), indicating that growth form represents a major axis of microbiome variation among these four octocoral hosts.

The mechanistic basis for these morphology-dependent microbial associations may be attributed to differential environmental exposure and microbial recruitment pathways. Encrusting species, which form mats directly over reef substrates, experience dual microbial recruitment from both the water column and benthic environments. As McCauley et al. (2016) demonstrated with *E. caribaeorum*, this close proximity to the substratum may facilitate infiltration of substrate-associated microbes into coral tissue, potentially expanding the available microbial pool. By contrast, the erect structures of branching octocorals limit microbial recruitment to the water column, resulting in more constrained communities. This pattern of morphological constraint on microbial diversity is consistent with findings from other benthic marine systems, including seaweeds (Lemay et al., 2021) and sponges (Gloeckner et al., 2014; Poppell et al., 2014). While relative abundance data provide valuable insights into community composition, measuring absolute microbial abundances (e.g., through quantitative methods such as flow cytometry; Props et al., 2017) would enhance interpretation of richness and evenness differences between species, as well as helping distinguish whether observed patterns reflect true shifts in bacterial loads or proportional changes within communities.

### Species-specific core microbiomes and *Endozoicomonas* dominance

Our core microbiome analysis, employing multiple prevalence thresholds (75%, 90%, and 100%), revealed species-specific structure that illuminate fundamental aspects of octocoral-bacteria associations.

We present 90% threshold results in the main text, validated through bootstrap resampling, with comparative analyses across all thresholds. This analysis confirmed the exceptional stability of identified core members across sampling scenarios, indicating these associations represent truly consistent host-microbe partnerships rather than sampling artifacts. At the 90% prevalence threshold, *Endozoicomonas* dominated the core microbiome of all four octocoral species (Figure 3). The ubiquity of *Endozoicomonas* across all examined species, combined with the presence of distinct ASV variants in each host, supports the hypothesis of species-specific coevolved symbioses within this bacterial genus. Despite harboring *Endozoicomonas*, *B. asbestinum* diverged from this pattern: an unclassified Gammaproteobacteria ASV constituted the most abundant core member (18.18%), slightly exceeding the combined *Endozoicomonas* abundance. This suggests that *B. asbestinum* employs a different microbial association strategy despite also harboring *Endozoicomonas* symbionts.

Systematic comparison across prevalence thresholds revealed contrasting patterns of core stability between encrusting and branching octocorals. *E. caribaeorum* exhibited the largest reduction in core size across thresholds, declining from 90 ASVs at 75% prevalence to 24 ASVs at 90%, and ultimately to 5 ASVs at 100% prevalence (see also supplementary material). This steep decline suggests higher inter-individual variation in bacterial colonization within encrusting species, potentially reflecting their greater exposure to benthic microbial sources. In contrast, branching species demonstrated remarkable threshold stability: *E. flexuosa* maintained identical cores at 90% and 100% thresholds, while *M. muricata* reduced minimally from 2 to 1 ASVs. This pattern of stability indicates a tighter host control over bacterial community assemblages in branching octocorals and potentially reflects vertical transmission or highly selective horizontal acquisition of *Endozoicomonas* symbionts. Additionally, shared ASVs emerged between encrusting species, including members of *Endozoicomonas,* Hyphomonadaceae, and Rhizobiaceae (Supplementary Table S4), further supporting morphology-based differentiation in bacterial associations.

*Endozoicomonas* performs several key functions in octocoral holobionts. Metabolic support through glucose production and protection against oxidative stress are likely provided by these symbionts to their octocoral hosts (Ding et al. 2016). Additionally, members of this genus possess chitinase and chitin-binding protein-encoding genes, enabling them to potentially hydrolyze this molecule present in octocoral tissues (da Silva *et al*. 2023), which may in turn facilitate nutrient recycling within the holobiont. The pronounced species-specificity and consistent prevalence of distinct *Endozoicomonas* lineages observed in our study reinforces emerging evidence for co-evolutionary relationships between these bacterial symbionts and their octocoral hosts (Bayer et al. 2013, La Rivière et al. 2013, Neave et al. 2017, Pogoreutz et al. 2018, Pollock et al. 2018). The phylogenetic constraint on *Endozoicomonas* associations implies either vertical transmission or highly specific horizontal acquisition mechanisms that maintain host fidelity over evolutionary timescales. Further research is needed to elucidate the specific functional roles of *Endozoicomonas* in western Atlantic octocorals and the mechanisms underlying their host specificity.

### Beyond Endozoicomonas: diverse bacterial associates in octocoral microbiomes

In addition to *Endozoicomonas*, several other bacterial groups showed strong morphology-specific associations. Some colonies of *E. flexuosa* and *M. muricata* were characterized by high abundances of *Mycoplasma*, a pattern consistent with previous findings in Caribbean octocorals where this genus comprised 16-36% of total bacterial reads. *Mycoplasma* is an abundant bacterial group in tropical octocorals (Shirur, Jackson and Goulet 2016; McCauley, Jackson and Goulet 2020; Monti *et al*. 2023; Reigel and Hellberg 2023) and temperate gorgonians (van de Water, Allemand and Ferrier-Pagès 2018). The functional role of *Mycoplasma* in octocorals remains poorly understood, though it has been previously linked to nutrient uptake from coral-captured prey, as it was shown to be located exclusively within nematocysts in the mucus layer of the scleractinian coral *Lophelia pertusa* (Neulinger et al., 2009). While *Mycoplasma* may act as a parasite in some animals, in temperate gorgonian corals it appears to be a commensal or mutualistic symbiont and is often a dominant member of the microbiome in healthy colonies (van de Water et al., 2018).

In contrast to the branching species, the encrusting octocorals harbored more taxonomically diverse bacterial communities with different dominant groups. *B. asbestinum* was characterized by high abundances of unclassified Gammaproteobacteria (18.18% mean relative abundance), which constituted the most abundant taxon in its microbiome. Gammaproteobacteria and Alphaproteobacteria are widely recognized as major constituents of octocoral microbial communities across diverse marine environments and have been shown to form stable associations with various coral species. Additionally, members of the Alphaproteobacteria, including Marinobacter species, may play important roles in holobiont health through pathogen inhibition (Alagely *et al*. 2011). Rhodobacteraceae, members of the Alphaproteobacteria class, were consistently present in *B. asbestinum* and have been documented in other octocoral species.

*E. caribaeorum* exhibited an even more complex bacterial assemblage, including diverse Alphaproteobacteria groups; the BD1-7 clade (a member of *Cellvibrionales*, family Spongiibacteraceae, that is commonly found in both Mediterranean and Caribbean gorgonians; van de Water et al., 2018; Monti et al., 2023), *Filomicrobium*, and notably, archaeal representatives including Candidatus *Nitrosopumilus*. The concurrent presence of ammonia-oxidizing *Candidatus Nitrosopumilus* and nitrogen-fixing *Rhizobiales* members suggests *E. caribaeorum* possesses integrated nitrogen cycling capacity, encompassing both nitrogen fixation and ammonia oxidation. This dual capability may provide enhanced nutrient acquisition efficiency, particularly valuable in the coastal waters adjacent to highly urbanized areas such as the reefs in Southeast Florida. *Rhizobiales* have also been reported as common associates in Indo-Pacific soft corals including *Litophyton* sp., where they contribute to nitrogen cycling alongside other metabolically diverse bacteria (Park et al., 2021), and in Caribbean octocorals including *Antillogorgia elisabethae*, where they occur as stable low-abundance symbionts (Galkiewicz & Kellogg, 2008).

Although assessing the health status of sampled octocorals was beyond the scope of this study, our collection protocol targeted visually healthy colonies, and the consistent presence of *Mycoplasma* in branching species, and Gammaproteobacteria in encrusting species, supports their potential roles as beneficial symbionts rather than pathogenic agents. This morphology-specific bacterial association, with *Mycoplasma* dominating branching species while Gammaproteobacteria and diverse Alphaproteobacteria groups characterized encrusting species, reinforces the link between octocoral growth form and microbial community structure. This pattern likely reflects combined differences in feeding strategies, environmental exposure, and host-microbe selection mechanisms between these morphologically distinct groups. Full understanding of specific bacterial symbioses within metazoan hosts remains difficult and nebulous, in spite of intensifying research. Almost every eukaryotic lineage has unique mechanisms for the initiation, retention, evolution for their symbiotic associations (Lajoie and Parfrey 2022). For example, some molluscan symbioses depend greatly on chemosynthetic bacteria for energy, while most marine sponge species can actively select the bacteria they filter or allow to pass through their choanocyte chambers (Ip *et al*. 2021; Díez-Vives *et al*. 2022). Stable symbiont communities can be viewed as traits for the holobiont, and in some instances act as drivers for host diversification (Mestre *et al*. 2020). At present, we do not know the full extent and how octocoral species depend on the bacterial symbionts characterized in this study, even though selective processes are likely at work. Lastly, the presence of potentially pathogenic taxa in a host without noticeable signs of disease points to a functionally operating immune system in the individual.

### Temporal stability and environmental resilience

This study revealed spatiotemporal stability in octocoral-associated bacterial communities over the three-year sampling period. Although spatial and temporal factors significantly affected total bacterial composition, they only explained 2% and 1.8% of the total bacterial variation, respectively (Table 3). Notably, *E. caribaeorum* exhibited a significant decrease in alpha diversity parameters between 2023 and 2025, and 2024 and 2025 sampling time points, while *M. muricata* showed significant differences in Simpson diversity only between 2023 and 2025 (Figure S5). Because the 2023 and 2024 sampling time points occurred during Florida’s wet season (July-September) and the 2025 sampling occurred during the dry season (January-February), these results may indicate that seasonal conditions may affect the number of bacterial taxa associated with *E. caribaeorum* and to some degree *M. muricata.*. Nonetheless, while spatiotemporal factors influenced octocoral bacterial composition to some degree, the communities remained largely stable overall. It is noteworthy that our temporal sampling comprised only three time points over three years, with uneven seasonal representation. More frequent sampling across multiple annual cycles would better resolve seasonal effects and inter-annual variability.

The temporal stability observed in most species supports the concept of octocoral holobionts as stable ecological units with conserved host-microbe associations. This likely reflects strong host control over microbial colonization and the establishment of resilient core communities that persist despite environmental fluctuations. Such stability may be particularly important for long-lived colonial organisms like octocorals that must maintain consistent physiological functions over extended periods.

## Conclusions

This study provides comprehensive insights into the bacterial communities associated with four ecologically important and widespread Florida octocoral species, revealing that host identity and morphology are the primary determinants of microbiome composition and diversity. The pronounced host-specific clustering of bacterial communities, coupled with the dominance of distinct *Endozoicomonas* lineages in each species, indicates long-term co-evolutionary relationships between octocorals and their microbial symbionts. The striking differences between encrusting and branching morphologies, with encrusting species exhibiting diverse, generalist microbiomes, and branching species maintaining specialized, low-diversity communities dominated by *Endozoicomonas* and *Mycoplasma*; highlight the importance of growth form in shaping microbial associations. These morphology-dependent patterns likely reflect differences in environmental exposure, microbial recruitment pathways, and feeding strategies between these morphologically distinct groups. The overall temporal and spatial stability of octocoral microbiomes across a three-year period and multiple reef sites underscores the resilience of these host-microbe partnerships and suggests strong host control mechanisms maintain consistent bacterial communities despite strong environmental variation. However, the seasonal shifts observed in some species indicate that octocoral microbiomes retain some flexibility to respond to environmental changes. As Florida’s coral reefs face increasing anthropogenic pressures and climate change impacts, understanding the ecological and functional significance of these stable yet flexible microbiomes will be crucial for predicting octocoral responses to future environmental challenges and developing effective conservation strategies for these foundational species on Caribbean reef ecosystems.

## Acknowledgements

This work was conducted under the State of Florida Special Activity License SAL-25-2666-SRP which authorized octocoral collection. We thank Paisley Samuel for her outstanding laboratory assistance, as well as Andrew Bauman, and Faith Borak, Nicholas Jones and Manuel Ploner for field assistance with sample collections.

## Author contributions

RL: Investigation, Formal Analysis, Data Curation, Visualization, Methodology, Writing - Original Draft, Resources. MGC: Formal Analysis, Writing - Review & Editing. JVL: Conceptualization, Supervision, Resources, Writing - Review & Editing.

## Data availability

All sequence data used in this study are available through under NCBI BioProject PRJNA1373357 (https://www.ncbi.nlm.nih.gov/bioproject/PRJNA1373357). R files and scripts used in this work can be found on GitHub (https://github. com/ ronen liberman/).

