## Supplementary material for "Host species identity drives temporally stable octocoral microbiomes in Florida": Table S1:S6; Figure S1:8

**Table S1. Geographic coordinates and depths of octocoral collection sites in southeastern Florida.** Samples were collected from shallow (5-8 m) and deep (15-20 m) reef sites in Miami-Dade and Broward counties during July 2023 - February 2025

| Site name | latitude | longtitude | Depth (m) |
|---|---|---|---|
| Miami shallow | 25.50566 | -80.0623 | 5-8 |
| Miami Deep | 25.5057 | -80.0527 | 15-18 |
| *Broward Shallow* | 26.1806 | -80.0410 | 5-18 |
| *Broward Deep* | 26.0891 | -80.0535 | 15-20 |

**Table S2. Summary of 16S rRNA gene sequencing results for octocoral-associated bacterial communities and seawater controls.** Total reads and amplicon sequence variants (ASVs) detected across four octocoral species (*Briareum asbestinum, Erythropodium caribaeorum, Eunicea flexuosa*, and *Muricea muricata*) and seawater samples collected from coral reef sites. Data represent post-quality filtering results following removal of chloroplast and mitochondrial sequences, chimeric reads, and low-quality sequences. Sequences were processed using QIIME2 and assigned to ASVs at 100% sequence similarity

| Group | Number of samples | Total Reads | Total ASVs |
|---|---|---|---|
| *E. caribaeorum* | 65 | 5,887,612 | 31,745 |
| *B. asbestinum* | 60 | 4,397,458 | 20,515 |
| *E. flexuosa* | 31 | 2,238,955 | 6,200 |
| *M. muricata* | 52 | 4,563,692 | 5,554 |
| Seawater | 20 | 1,836,298 | 5,263 |

**Table S3. Sensu stricto core bacterial taxa in Florida octocoral microbiomes.** Complete list of bacterial ASVs defined as taxa present in 100% of samples within each octocoral species. Each Table shows genus-level taxonomic identification and mean relative abundance. Genus ID refers to the global systematic numbering of ASVs within each bacterial genus, ensuring consistent identification across species. Bold font indicates unique ASVs that were both statistically significant in ANCOMBC-BC2 differential abundance testing and identified as core microbiome members. Statistical significance from ANCOMBC-BC2 analysis is indicated as: *** $p < 0.001$ and * $p < 0.05$.

| Species | Genus ID | Mean abundance (%) |
|---|---|---|
| ***B. asbestinum*** | ***Endozoicomonas*_4** | **9.5690*** |
| *B. asbestinum* | unclassified Rhodobacteraceae_1 | 1.1590 |
| *E. caribaeorum* | *Filomicrobium*_1 | 0.7540 |
| *E. caribaeorum* | *Filomicrobium*_2 | 0.2460 |
| ***E. caribaeorum*** | **unclassified Alphaproteobacteria_5** | **3.8130*** ** |
| *E. caribaeorum* | unclassified Rhizobiaceae_1 | 0.2330 |

| | | |
|---|---|---|
| *E. caribaeorum* | unclassified Rhodobacteraceae_1 | 1.1730 |
| *E. flexuosa* | *Endozoicomonas*_6 | 7.4190 |
| *M. muricata* | *Endozoicomonas*_5 | 7.7180 |

Table S4. ASV shared between host species and their taxonomic details

| Core threshold | #ASVs shared between species | Host species identity | ASV associated bacterial order | ASV associated bacterial family | ASV associated bacterial genus |
|---|---|---|---|---|---|
| 100% | 2 | *B. asbestinum; E. caribaeorum* | Rhodobacterales | Rhodobacteraceae | unclassified Rhodobacteraceae |
| 90% | 2 | *B. asbestinum; E. caribaeorum* | Oceanospirillale s | Endozoicomonadaceae | Endozoicomonas |
| 90% | 2 | *B. asbestinum; E. caribaeorum* | Caulobacterales | Hyphomonadaceae | unclassified Hyphomonadaceae |
| 90% | 2 | *B. asbestinum; E. caribaeorum* | Rhizobiales | Rhizobiaceae | unclassified Rhizobiaceae |
| 90% | 2 | *B. asbestinum; E. caribaeorum* | Rhodobacterales | Rhodobacteraceae | unclassified Rhodobacteraceae |
| 75% | 3 | *B. asbestinum; E. caribaeorum; E. flexuosa* | Rhodobacterales | Rhodobacteraceae | unclassified Rhodobacteraceae |
| 75% | 2 | *B. asbestinum; E. caribaeorum* | Oceanospirillale s | Endozoicomonadaceae | Endozoicomonas |

| | | | | | |
|---|---|---|---|---|---|
| 75% | 2 | *B. asbestinum; E. caribaeorum* | Rhizobiales | Hyphomicrobiaceae | Filomicrobium |
| 75% | 2 | *B. asbestinum; E. caribaeorum* | Caulobacterales | Hyphomonadaceae | unclassified Hyphomonadaceae |
| 75% | 2 | *B. asbestinum; E. caribaeorum* | Rhizobiales | Methyloligellaceae | Methyloceanibacter |
| 75% | 2 | *B. asbestinum; E. caribaeorum* | Microtrichales | Microtrichaceae | Sva0996_marine_group |
| 75% | 2 | *B. asbestinum; E. caribaeorum* | Rhizobiales | Rhizobiaceae | unclassified Rhizobiaceae |
| 75% | 2 | *B. asbestinum; E. caribaeorum* | Vibrionales | Vibrionaceae | Photobacterium |

**Table S5. Individual species beta diversity analysis** showing PERMANOVA results for temporal and spatial effects on octocoral-associated microbial communities. F-statistics, P-values, $R^2$ values, and sample sizes (N) are presented for each species and factor combination. Temporal factor represents year-to-year variation, while spatial factor represents site-to-site variation within each octocoral species.

| Factor | Species | F-statistic | P-value | $R^2$ | N |
|---|---|---|---|---|---|
| Temporal | *B. asbestinum* | 1.920 | 0.005 | 0.084 | 45 |
| Temporal | *E. flexuosa* | 1.862 | 0.008 | 0.134 | 27 |
| Temporal | *E. caribaeorum* | 1.822 | 0.001 | 0.065 | 55 |
| Temporal | *M. muricata* | 2.120 | 0.002 | 0.081 | 51 |
| Spatial | *B. asbestinum* | 2.554 | 0.001 | 0.157 | 45 |
| Spatial | *E. flexuosa* | 1.541 | 0.029 | 0.167 | 27 |

| Factor | Species | F-statistic | P-value | $R^2$ | N |
|---|---|---|---|---|---|
| Spatial | *E. caribaeorum* | 1.658 | 0.001 | 0.089 | 55 |
| Spatial | *M. muricata* | 1.971 | 0.003 | 0.112 | 51 |

**Table S6. Pairwise comparisons of bacterial community composition between octocoral species and seawater controls.** Results show PERMANOVA F-statistics, explained variance ($R^2$), raw p-values, and multiple comparison-corrected p-values (Bonferroni) for bacterial communities based on Bray-Curtis dissimilarity. Sample sizes: seawater (n =15), *Briareum asbestinum* (n = 45), *Eunicea flexuosa* (n = 27), *Erythropodium caribaeorum* (n = 55), *Muricea muricata* (n = 51). Sequences were rarefied to 20,000 reads per sample prior to analysis.

| Group | F-statistic[1] | $R^2$[2] | P-value | P. adjusted (Bonferroni) |
|---|---|---|---|---|
| Seawater vs *B. asbestinum* | 23.470 | 0.288 | 0.001 | 0.01 |
| Seawater vs *E. flexuosa* | 13.432 | 0.251 | 0.001 | 0.01 |
| Seawater vs *E. caribaeorum* | 27.525 | 0.288 | 0.001 | 0.01 |
| Seawater vs *M. muricata* | 23.307 | 0.267 | 0.001 | 0.01 |
| *B. asbestinum* vs *E. flexuosa* | 13.492 | 0.162 | 0.001 | 0.01 |
| *B. asbestinum* vs *E. caribaeorum* | 24.870 | 0.202 | 0.001 | 0.01 |
| *B. asbestinum* vs *M. muricata* | 28.613 | 0.233 | 0.001 | 0.01 |
| *E. flexuosa* vs *E. caribaeorum* | 16.008 | 0.167 | 0.001 | 0.01 |
| *E. flexuosa* vs *M. muricata* | 12.953 | 0.146 | 0.001 | 0.01 |
| *E. caribaeorum* vs *M. muricata* | 35.056 | 0.252 | 0.001 | 0.01 |

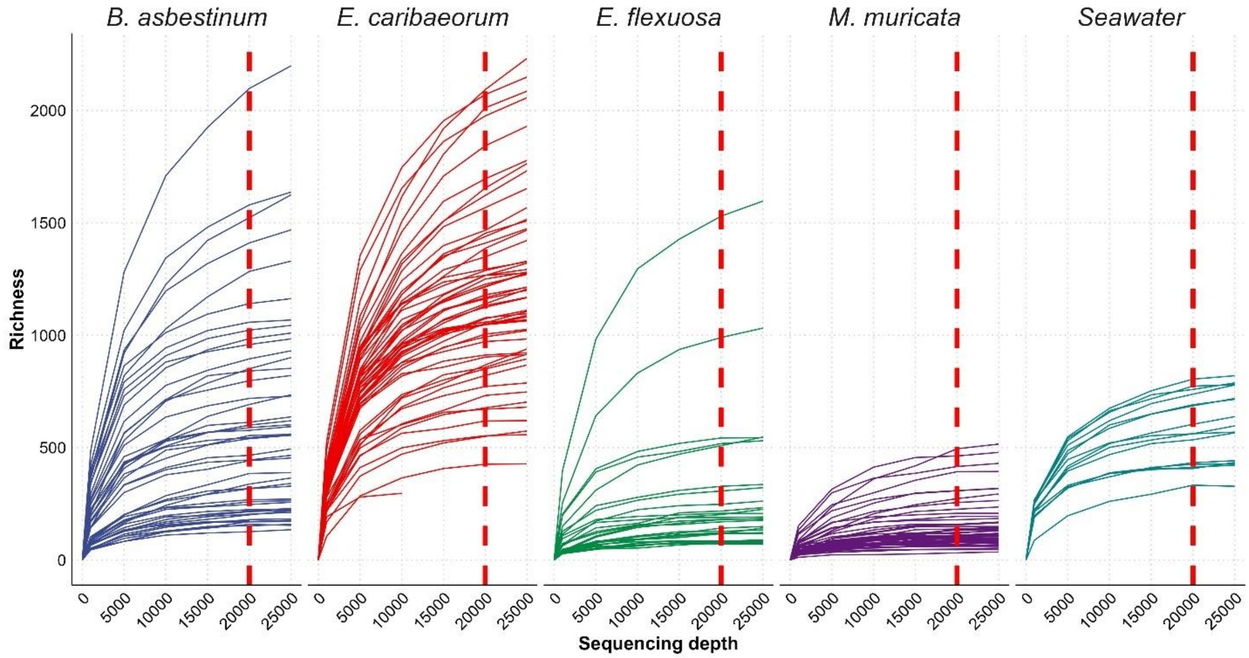


**Figure S1. Rarefaction curves showing bacterial community richness across sampling sequencing depth for octocoral species and seawater controls.** Individual curves represent bacterial ASV richness for each sample as a function of sequencing depth for *Briareum asbestinum* (blue), *Erythropodium caribaeorum* (red), *Eunicea flexuosa* (green), *Muricea muricata* (purple), and seawater (teal). The red dashed vertical line indicates the rarefaction depth of 20,000 reads per sample used for downstream analyses. Curves demonstrate adequate sampling depth for capturing bacterial diversity across all sample types, with most samples approaching richness saturation at the selected rarefaction threshold.

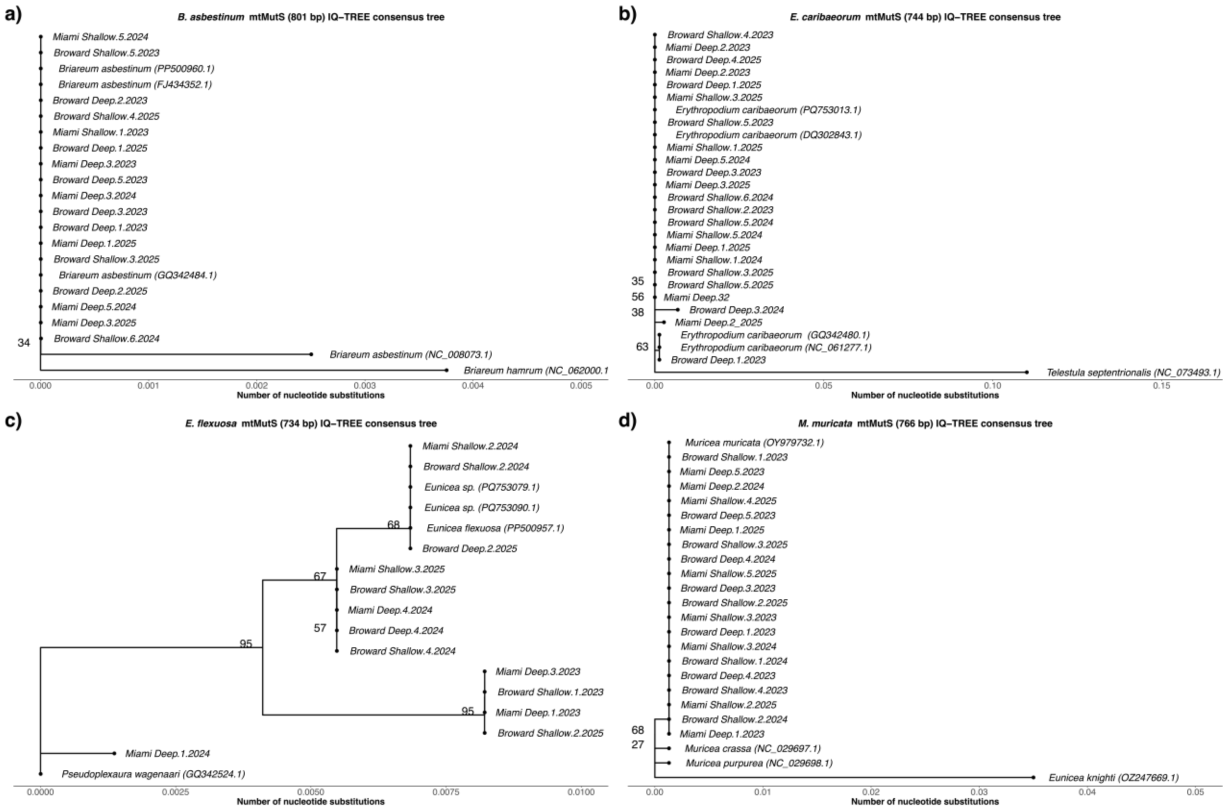


**Figure S2.** Individual octocoral phylogenetic trees inferred from the *mtMuts* gene for species a) *Briareum asbestinum*, b) *Erythropodium caribaeorum*, c) *Eunicea flexuosa*, and d) *Muricea muricata*. Topologies were inferred by maximum likelihood (ML) in IQ-TREE. Node labels indicate ML bootstrap support values. Each tree was rooted to an outgroup sequence downloaded form Genebank, placed at the base of the phylogeny. Additional sequences from GenBank were included for phylogenetic context for each species. All sequences from GenBank are shown in parentheses.

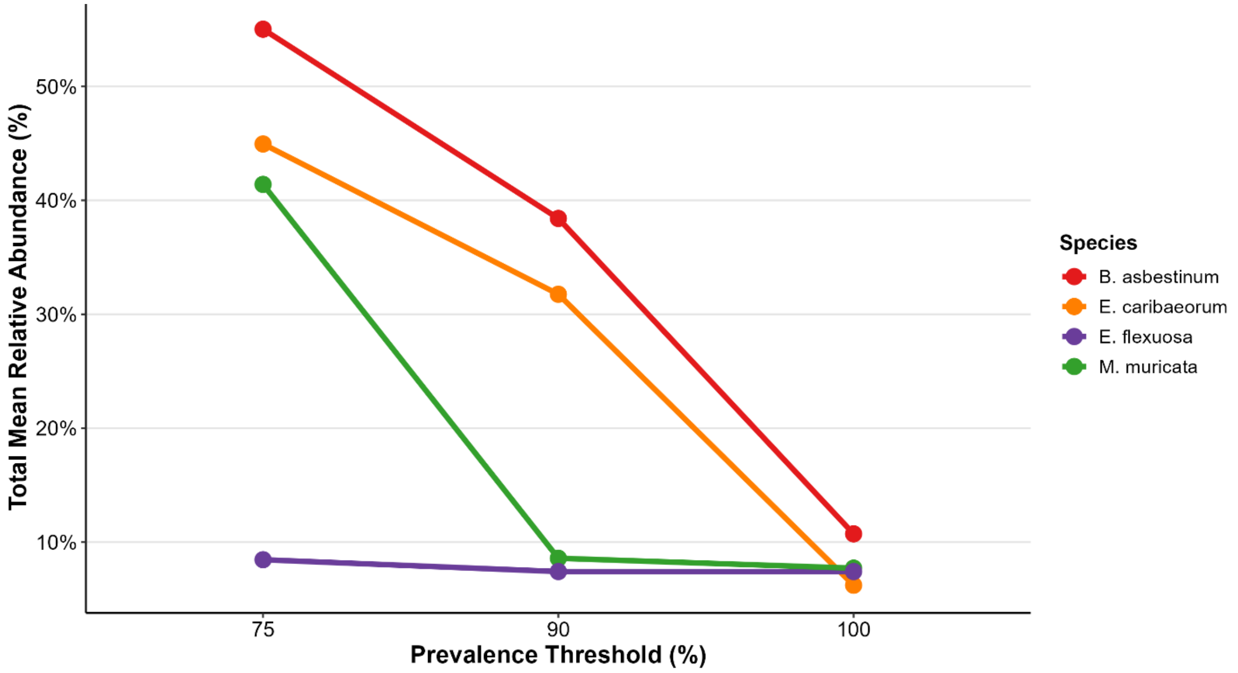


**Figure S3**. Cumulative mean relative abundance of core bacterial taxa across prevalence thresholds in four octocoral species. Line plots show the total proportion of the microbiome represented by core ASVs at three prevalence thresholds (75%, 90%, and 100%) for *Briareum asbestinum* (red), *Erythopodium caribaeorum* (orange), *Eunicea flexuosa* (purple), and *Muricea muricata* (green).

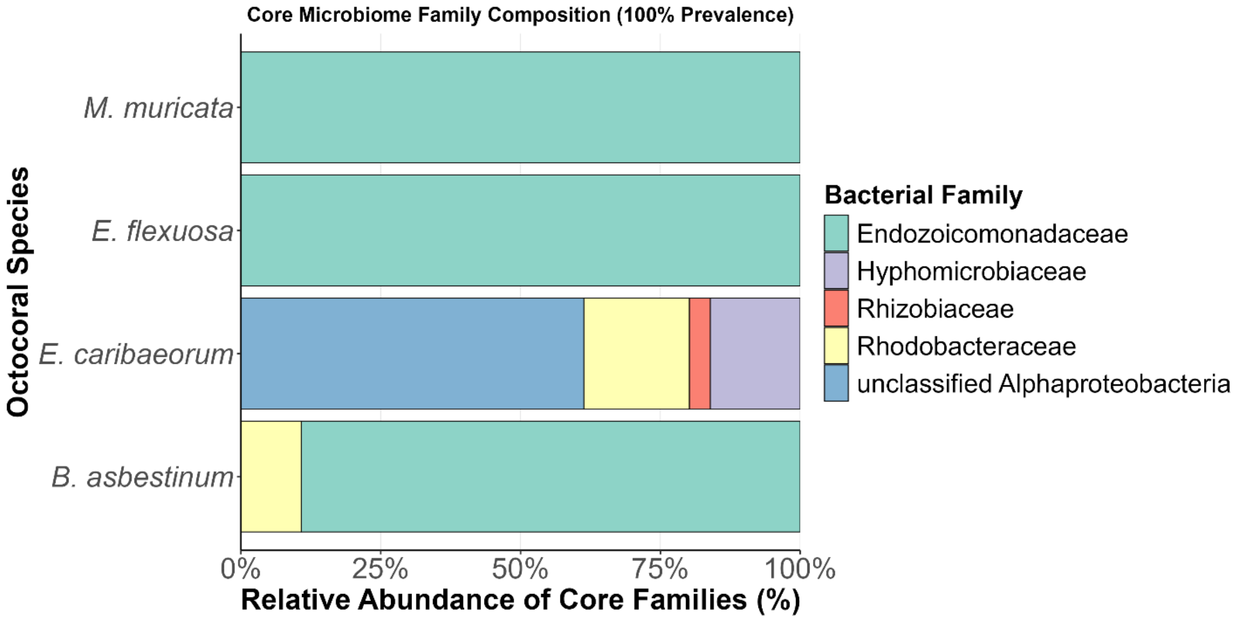


**Figure S4. Bacterial family composition of sensu stricto core microbiomes in four octocoral species.** Stacked horizontal bars show the relative abundance of bacterial families within the core microbiome defined at 100% prevalence (present in all samples) for each species.

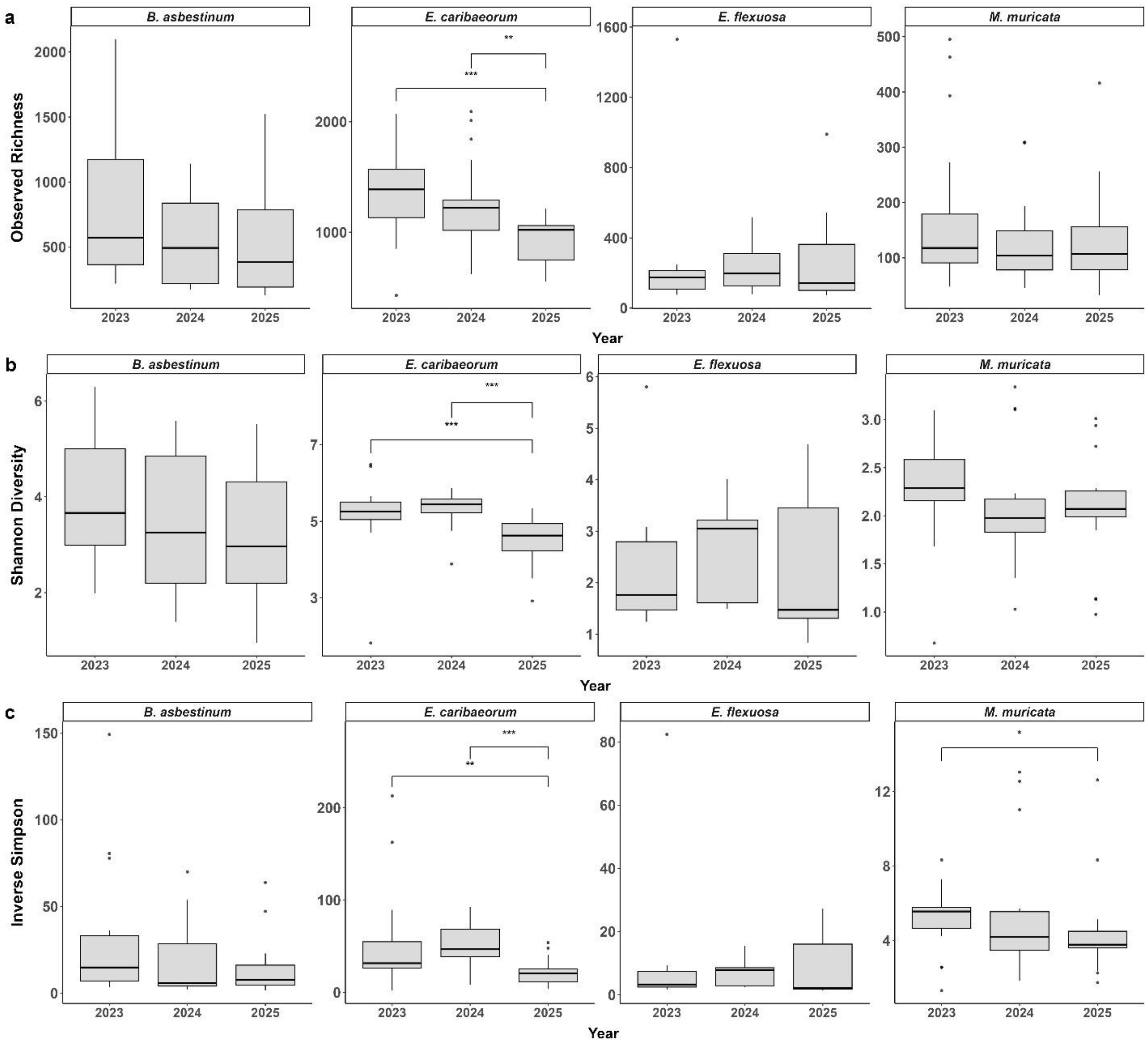


**Figure S5. Temporal changes in alpha diversity metrics of bacterial communities within octocoral species.** Boxplots show (a) observed richness, (b) Shannon diversity index, and (c) inverse Simpson diversity index for bacterial communities associated with *Briareum asbestinum, Erythopodium caribaeorum, Muricea muricata, and Eunicea flexuosa* across sampling years. Data represents all samples collected from Broward and Miami counties from July 2023 to February 2025. Sequences were rarefied to 20,000 reads per sample to normalize sequencing depth. Box plots display median (center line), first and third quartiles (box boundaries), and 1.5× interquartile range (whiskers), with outliers shown as individual points. Significance brackets with asterisks indicate pairwise differences between years within each species (Wilcoxon rank-sum test: * $p < 0.05$, ** $p < 0.01$, *** $p < 0.001$).

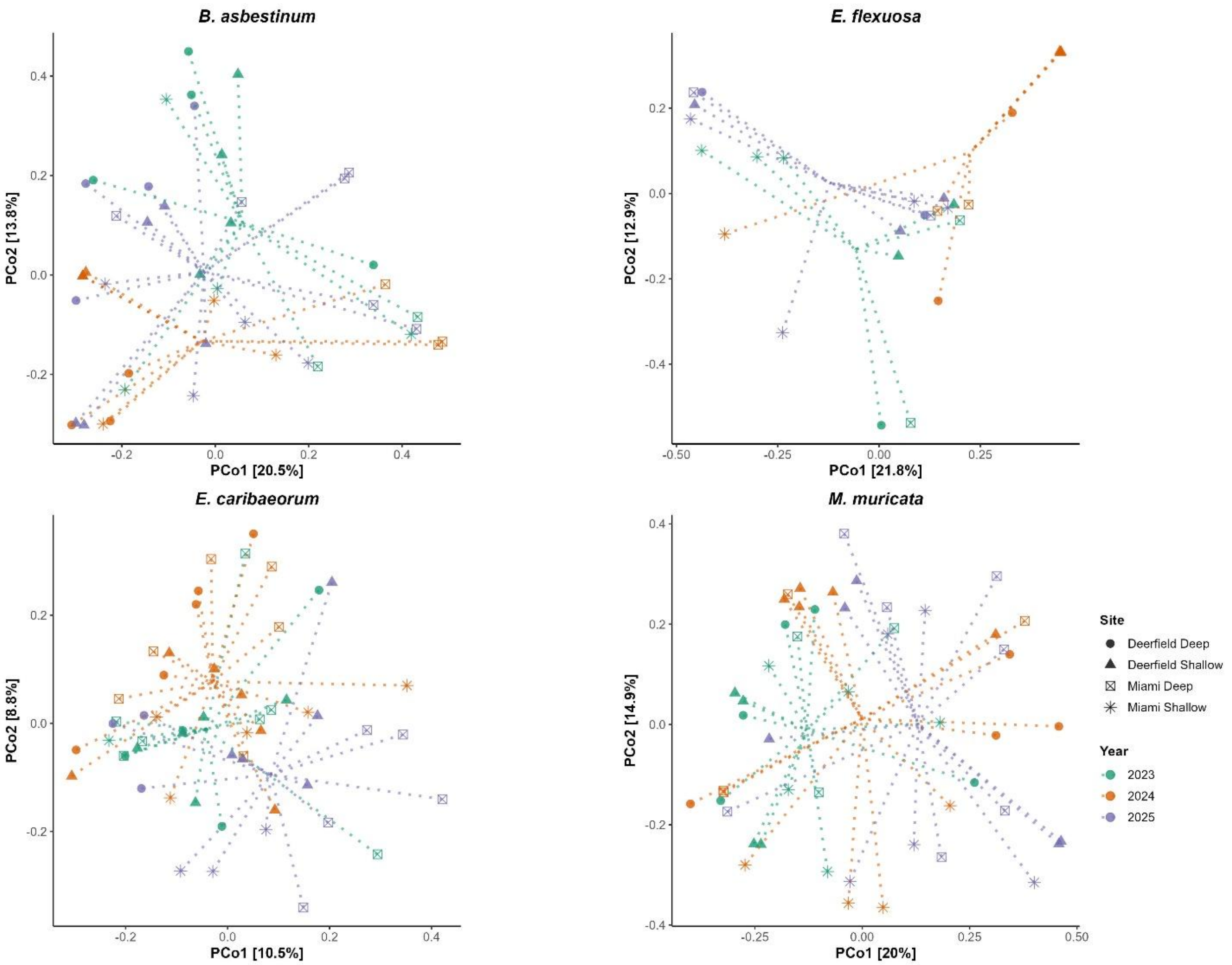


**Figure S6. Principal coordinate analysis (PCoA) ordinations showing temporal variation in octocoral-associated bacterial communities.** Each panel represents a different octocoral species with samples colored by collection year (2023, 2024, 2025) and shaped by sampling sites. Ordinations are based on Bray-Curtis dissimilarity matrices of bacterial community composition. Centroids represent the average community composition for each year within each species.

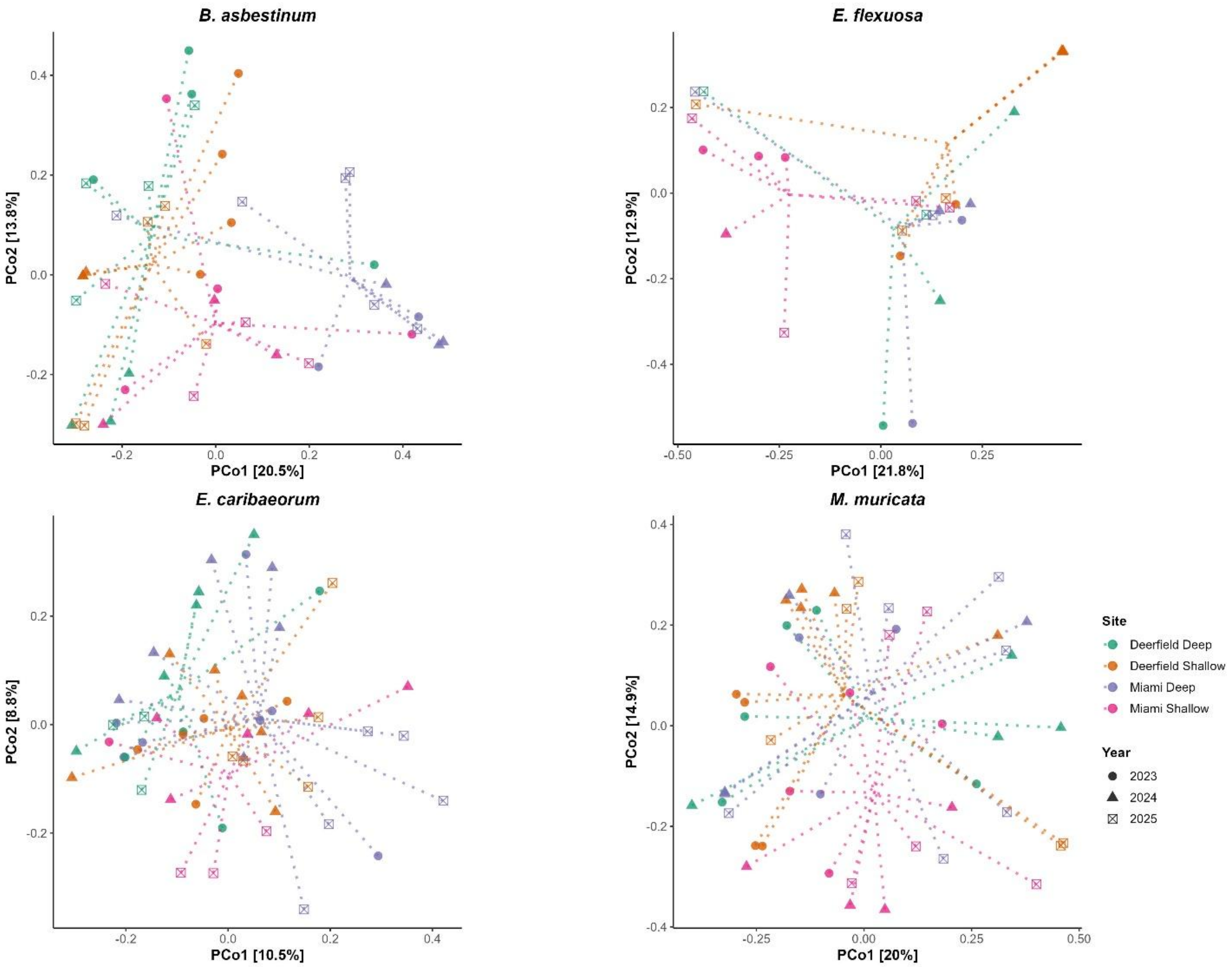


**Figure S7. Principal coordinate analysis (PCoA) ordinations showing spatial variation in octocoral-associated bacterial communities.** Each panel represents a different octocoral species with samples colored by sampling site and shaped by collection year. Sites include shallow (5 – 10 m) and deep (15 - 20 m) locations at Deerfield Beach and Miami. Ordinations are based on Bray-Curtis dissimilarity matrices of bacterial community composition. Centroids represent the average community composition for each site within each species.

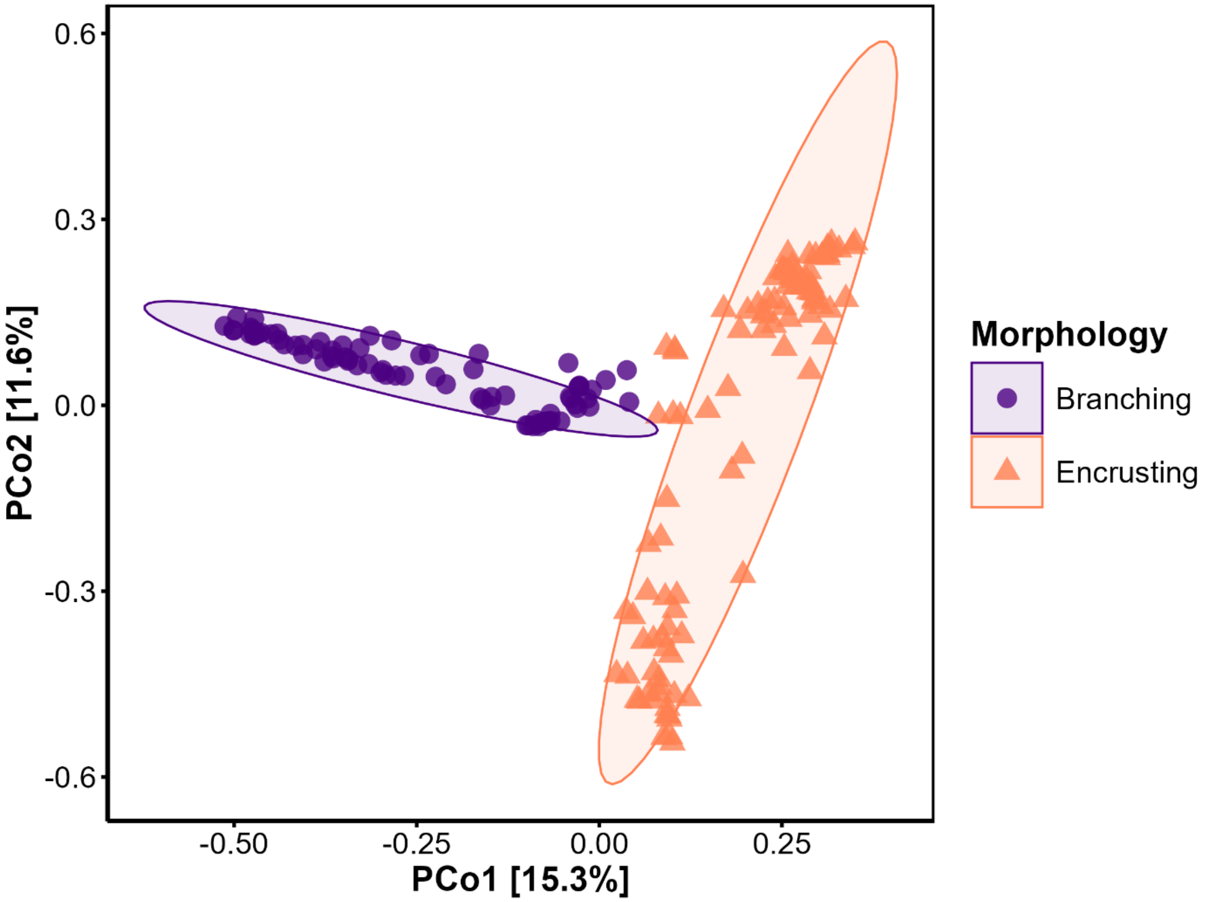


**Figure S8. Morphological differences in octocoral bacterial community composition.** Principal coordinate analysis (PCoA) based on Bray-Curtis dissimilarity demonstrates distinct clustering of bacterial communities by colony morphology (branching vs. encrusting). Data represents rarefied bacterial ASV abundance (20,000 reads per sample) from branching species (*Eunicea. flexuosa and Muricea muricata*) and encrusting species (*Briareum asbestinum and Erythopodium caribaeorum*), collected from southeastern Florida reefs. The first two axes explain 26.9% of total variation (PCo1: 15.3%, PCo2: 11.6%). Ellipses indicate 95% confidence intervals around morphology group centroids.
